# Naked Mole-Rat Hematopoietic Stem and Progenitors are Highly Quiescent with an Inherent Myeloid Bias

**DOI:** 10.1101/2021.08.01.454652

**Authors:** Stephan Emmrich, Alexandre Trapp, Frances Tolibzoda Zakusilo, Marco Mariotti, Maggie E. Straight, Zhihui Zhang, Spencer Gray, Michael G. Drage, Zhonghe Ke, Masaki Takasugi, Jan-Henning Klusmann, Vadim N. Gladyshev, Andrei Seluanov, Vera Gorbunova

## Abstract

Naked mole-rats (NMRs) are the longest-lived rodents yet their stem cell characteristics remain enigmatic. Here we comprehensively mapped the NMR hematopoietic landscape and identified unique features likely contributing to longevity. Adult NMRs form red blood cells in spleen and marrow, which is a neotenic trait. A myeloid bias towards granulopoiesis in concert with decreased B-lymphopoiesis defines the marrow composition, resembling fetal leukopoiesis. Very similar to primates, the primitive stem cell compartment is marked by CD34 and THY1. Remarkably, stem and progenitor respiration rates are as low as in human cells, while NMR cells show a strong expression signature for fatty acid metabolism. The pool of quiescent stem cells is higher than in mice, and the cell cycle of hematopoietic cells is prolonged. Our work provides a platform to study immunology and stem cell biology in an animal model of exceptional longevity.

**Teaser:** Juvenile features of hematopoiesis shape the blood system of the longest-lived rodent.

## Introduction

The naked mole-rat (*Heterocephalus glaber*) emerged as an animal model of exceptional longevity and resistance to age-associated diseases (*1*). At the size of a mouse these rodents reach a lifespan of over 30 years in captivity and do not display increased mortality with aging (*2*). NMR cells feature higher translation fidelity due to split processing of 28S rRNA (*3*), express a unique splicing product from the senescence- inducing INK4/ARF locus and maintain ample high molecular weight hyaluronic acid (HMW-HA) responsible for the resistance of NMRs to solid tumors (*4, 5*).

The blood is the most regenerative tissue, producing >10^14^ cells per year in humans (*6*). Fostered by recent advances in single-cell transcriptomics (*7*), hematopoiesis is viewed as a continuum of individual cells that traverse the differentiation process from unprimed hematopoietic stem and progenitor cells (HSPCs) directly into unipotent progenitors (*8*). Studies of the unperturbed hematopoietic system are most advanced in their understanding of HSPC hierarchies and concepts of stemness in mice (*9*). There are, however, fundamental differences in certain aspects of the blood system between mice and humans (*10*). At the genetic level, orthologs for one of the major murine HSC markers, Sca-1 (Ly6a), are found only in rodents but not in primates, carnivores, birds or fish. Interestingly, NMRs are among the few rodents without a Sca-1 ortholog (Table S1).

We developed a flow cytometry (FACS) labelling strategy using cross-reactive antibodies to sort, culture and transplant NMR hematopoietic stem and progenitor cells (HSPCs). A panel of six surface markers allowed to purify primitive stem cells with multi-lineage potential, distinct cell stages during early erythroid and T-lymphoid commitment (*11*) and distinguished the major blood leukocyte fractions. NMR HSPCs showed striking similarities to human HSPCs, such as a CD34^+^ compartment harboring primitive progenitors, marrow granulopoiesis and slow cell metabolism. Further adaptations involved a prolonged cell cycle duration, splenic erythropoiesis and retention of juvenile platelet and leukocyte counts in aged animals, revealing systemic deviations from traditional concepts of hematopoiesis to concertedly promote longevity. Our findings provide a comprehensive resource for the studies of immunosenescence, ‘inflammaging’ and stem cell biology in the naked mole-rat as a model of exceptional longevity.

## Results

### The developmental landscape of naked mole-rat hematopoiesis

To separate NMR hematopoietic cells, we screened 101 commercially available monoclonal antibodies (moAb; Table S7) against human, mouse, rat and guinea pig CD markers, and identified human CD11b, CD18, CD34 and CD90, mouse CD11b and CD125, rat Thy1.1 and guinea pig CD45 as cross-reactive to bind distinct subsets of viable NMR bone marrow (BM) cells (Fig. 1a). Antibodies were validated by isotypes, Fc blockers and cross-species comparison (Fig. S1a-b).

**Fig. 1.**
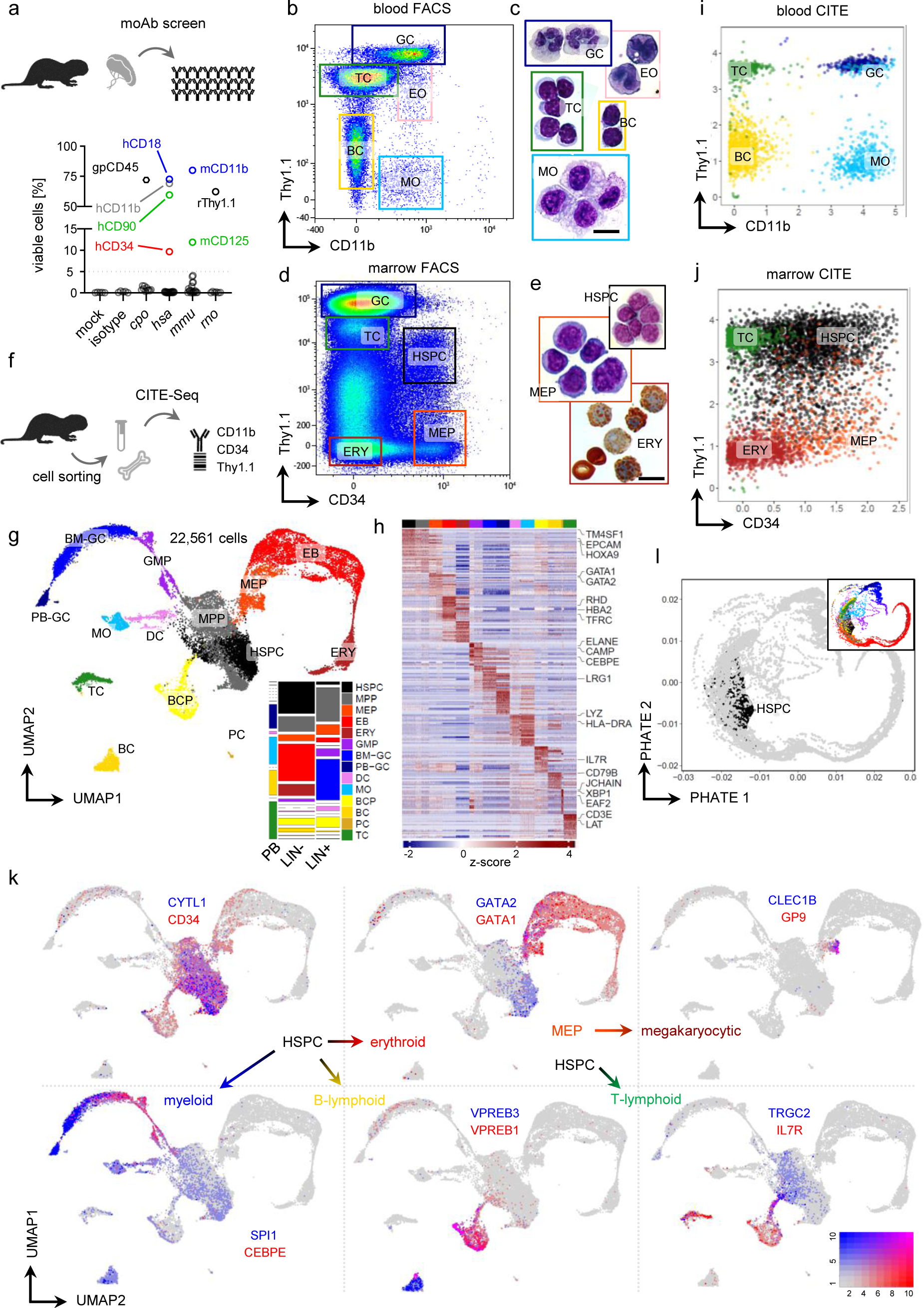
Purification of blood cell types and the developmental hierarchy in the marrow. **a**, Frequency of naked mole-rat bone marrow cells stained with cross-reactive antibodies (n=101); mock, unstained; *cpo*, guinea pig target host; *hsa*, human; *mmu*, mouse; *rno*, rat. Dotted line, 5% threshold unspecific binding. Representative FACS gating of **b**, blood (PB) stained with Thy1.1 and CD11b or **d**, marrow (BM) stained with Thy1.1 and CD34. Sorting gates: GC, neutrophil granulocytes; BC, B cells; TC, T cells; MO, monocytes; EO, eosinophils; HSPC, hematopoietic stem and progenitor cells; MEP, megakaryocytic erythroid progenitor; ERY, erythroid cells. May-Grünwald-Giemsa staining of sorted **c**, PB or **e**, BM cells; Scale bar 20µm, same magnification for each micrograph. **f**, Sorted PB (n=1) and BM (n=3) were used for CITE-Seq (G-L) with antibodies from (B-E). **g**, UMAP of Louvain-clustered single-cell transcriptomes, color legend is used throughout this dataset; 1799 differentially expressed genes were used for fGSEA-based cell type annotation. Tile-stack inset reflects relative cluster frequencies [y] and tissue library fractions of the dataset [x] as probability. MPP, multipotent progenitor; EB, erythroblast; GMP, granulocytic monocytic progenitor; BM-GC, marrow neutrophils; PB-GC, blood neutrophils; DC, dendritic cells; BCP, B cell progenitor; PC, plasma cells. **h**, Heatmap showing top 25 overexpressed genes by fold-change of 14 single-cell clusters from sorted BM and PB randomly downsampled to ≤ 500 cells, curated cell type markers are labelled. Scaled CITE-UMI counts per cell as i, Thy1.1 vs CD11b for PB and j, Thy1.1 vs CD34 for BM. k, UMAP-based Blendplots showing pairs of differentially expressed lineage markers conserved across species; gene1 (red, high expression), gene2 (blue, high expression) and co-expressing cells (purple). See scale on the right; expression, scaled UMI counts. l, PHATE model of single-cell transcriptomes, HSPC cluster is highlighted in black; inset depicts model colored by annotation from g, showing position of progeny cell types relative to HSPCs.

When we stained red blood cell (RBC)-depleted NMR peripheral blood (PB) with Thy1.1 and CD11b, the 5 major blood cell types could be distinguished: neutrophil granulocytes (GC), eosinophil granulocytes (EO), T cells (TC), B cells (BC) and monocytes (MO) (Fig. 1b). Cytochemistry of FACS-purified cells revealed multi-lobulated nuclei and a pH- neutral cytoplasm for Thy1.1hi/CD11b+ GCs and bi-lobulated nuclei with a high density of acidic granulation for Thy1.1int/CD11b+ EOs, with both populations exhibiting granulocytic scatter properties (Fig. 1c, Fig. S1c). Two populations with lymphocytic morphology and size are labelled as Thy1.1int/CD11b– and Thy1.1lo/CD11b–, while Thy1.1–/CD11b+ cells resemble MOs. BM labelling showed a distinct CD34hi/Thy1.1int population of monomorphic cells with small cytoplasm that was absent in PB. CD34 and CD90/THY1 are human stem cell markers (*12*); we thus hypothesized these cells to contain the HSPC compartment (Fig. 1d-e).

To increase cell type resolution we performed CITE-Seq (*13*) for CD11b, CD34 and Thy1.1 on sorted NMR PB and BM cell populations (Fig. 1f, Fig. S1d-e). A de novo transcriptome assembly from deep sampling of NMR whole marrow was prepared according to the FRAMA pipeline (*14*) and used for transcript annotation, which revealed hundreds of previously unannotated genes as well as thousands of novel transcript isoforms (Table S1, Fig. S1f-g). We referenced quality control data, clustering, cell cycle scoring and cell type annotation with a human and a murine droplet-based single-cell RNA-Sequencing (scRNA-Seq) datasets of hematopoietic hierarchies from the literature (Fig. S2-4, Table S2). Mapping of 11920 NMR orthologs to 22561 cells yielded 14 clusters which displayed a densely interconnected map of hematopoietic development (Fig. 1g). NMR hematopoietic cell types expressed canonical lineage markers and are associated with corresponding gene signatures in gene set enrichment analysis (GSEA; Fig. 1h, Table S2). The PB or BM fraction CITE counts each confirmed their FACS erythroid progenitor (MEP) cluster overexpressing both GATA1 and GATA2, maintaining CD34 and downregulating Thy1.1 levels. While MEPs resemble progenitor morphology, CD34– ERY cells mostly contain RBCs and reticulocytes (Fig. 1d-e). Canonical cell cycle marker expression revealed the HSPC cluster almost exclusively in G1, matching earlier findings of fate decisions uncoupled from cell division in mice (*15*). By contrast, multipotent progenitor (MPP), MEP, erythroblast (EB), B cell progenitor (BCP) and granulocyte monocyte progenitor (GMP) clusters have the highest G2/M and S phase signatures (Fig. S1h).

To confirm our annotations and to profile expression kinetics across cells and clusters we highlighted differentially expressed genes with conserved roles during hematopoiesis (*16*). Blended transcript expression showed strict confinement of NMR CYTL1, a bone mass modulator exclusively induced in human CD34+ HSPCs (*17*), to HSPC/MPP clusters, whereas CD34 is also expressed in MEPs and BCPs (Fig. 1k). Across humans, mouse and NMR HOXA9 expression was found in the most primitive HSPC cluster, with TM4SF1 specific for NMR HSPCs, whereas TM6SF1 as a marker of lymphomyeloid differentiation is conserved in rodents (Fig. S1i, 2f, 3f). Midkine (MDK), a pleitropic growth factor, emerged as a specific marker of NMR MEPs. Similar to human and mice however, NMR GATA1 determined erythroid commitment by specific expression in MEPs, EBs and erythroid cells (ERY), while GATA2 was expressed in HSCs and an MPP subset to merge with GATA1 in MEPs and early EBs (Fig. 1k, Fig. S2e, 3e). Interestingly, GP9, expressed at the surface of platelets, was specifically produced in MEP cells not expressing GATA2. This suggests NMR MEPs differentiate through a GATA2-dependent switch between the erythroid route along GATA1+/GATA2+/EPOR+ vs the GATA1+/GATA2–/GP9+/CLEC1B+ axis towards megakaryoblasts. In the lymphoid branch we found three developmental stages of CD20+ B cell clusters, with BCP showing exclusive expression of VPREB1, located on proB and preB cells (Fig. 1k). Cluster BC maintained VPREB3 overexpression, suggesting broad species conservation of successive VPREB gene waves as required for BC development. PU.1 (SPI1) is a transcription factor of the myeloid lineage with a role in HSC maintenance (*18*). Likewise, SPI1 was induced in naked-mole-rat MPPs in close proximity to GMPs (Fig. 1g-k), its absence in MEPs suggests conservation of the classic GATA1-PU.1 bi-modal switch. Thus SPI1 expression in the HSPC compartments marks the onset of myeloid commitment, converging with CEBPE into the granulocytic lineage, a pattern conserved in mouse (Fig. S3e). Strikingly, an clustering (*19*), projected HSC/HSPC clusters as central hubs connecting between clusters of the three major lineages (erythroid, lymphoid, myeloid) throughout every species and dataset we tested (Fig. 1l, Fig. S2g, 3g).

**Fig. 2.**
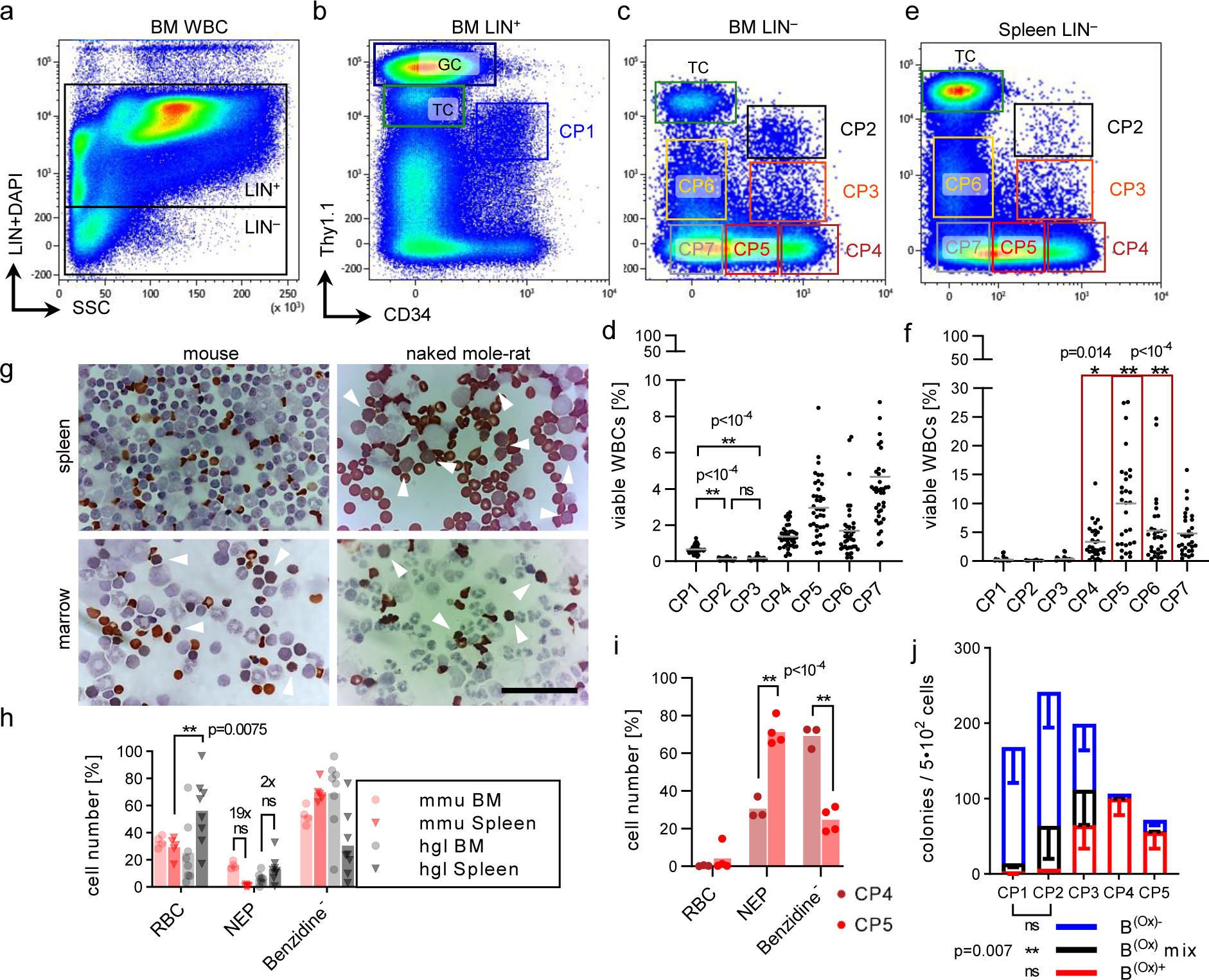
Normal erythropoiesis predominantly occurs in the spleen. Sorting strategy for the HSPC compartment with **a**, lineage (LIN = CD11b/CD18/CD90/CD125) depletion, **b**, gating of LIN^+^ CP1 and **c**, gating of LIN^−^ CP2-7. **d**, Frequencies of BM CP cell fractions. p-value determined by Brown-Forsythe’s One-way ANOVA; n=39; animal age range 1- 4yr. **e**, Representative gating of LIN^−^ CP2-7 in spleen. **f**, Frequencies of spleen CP cell fractions; n=30. p-value determined by Sidak’s Two-way ANOVA comparing BM vs spleen. **g**, Benzidine staining of whole spleen [top] or marrow [bottom] from 3 month old mice [left] or 3 year old naked mole-rats [right]. Scale bar 250µm, arrows indicate nucleated erythroid progenitors (NEPs). **h**, Relative counts of Benzidine-stained cytospins from whole spleen or WBM. p-value determined by Sidak’s Two-way ANOVA comparing BM vs spleen between mouse (n=5) and naked mole-rat (n=8). **i**, Relative counts of Benzidine-stained cytospins from naked mole-rat sorted spleen fractions. p-value determined by Sidak’s Two-way ANOVA comparing BM vs spleen; n=4. **j**, Benzidine-stained colony assays from sorted BM cells, n=3. Error bars denote s.d., p-value determined by Sidak’s Two-way ANOVA.

**Fig. 3.**
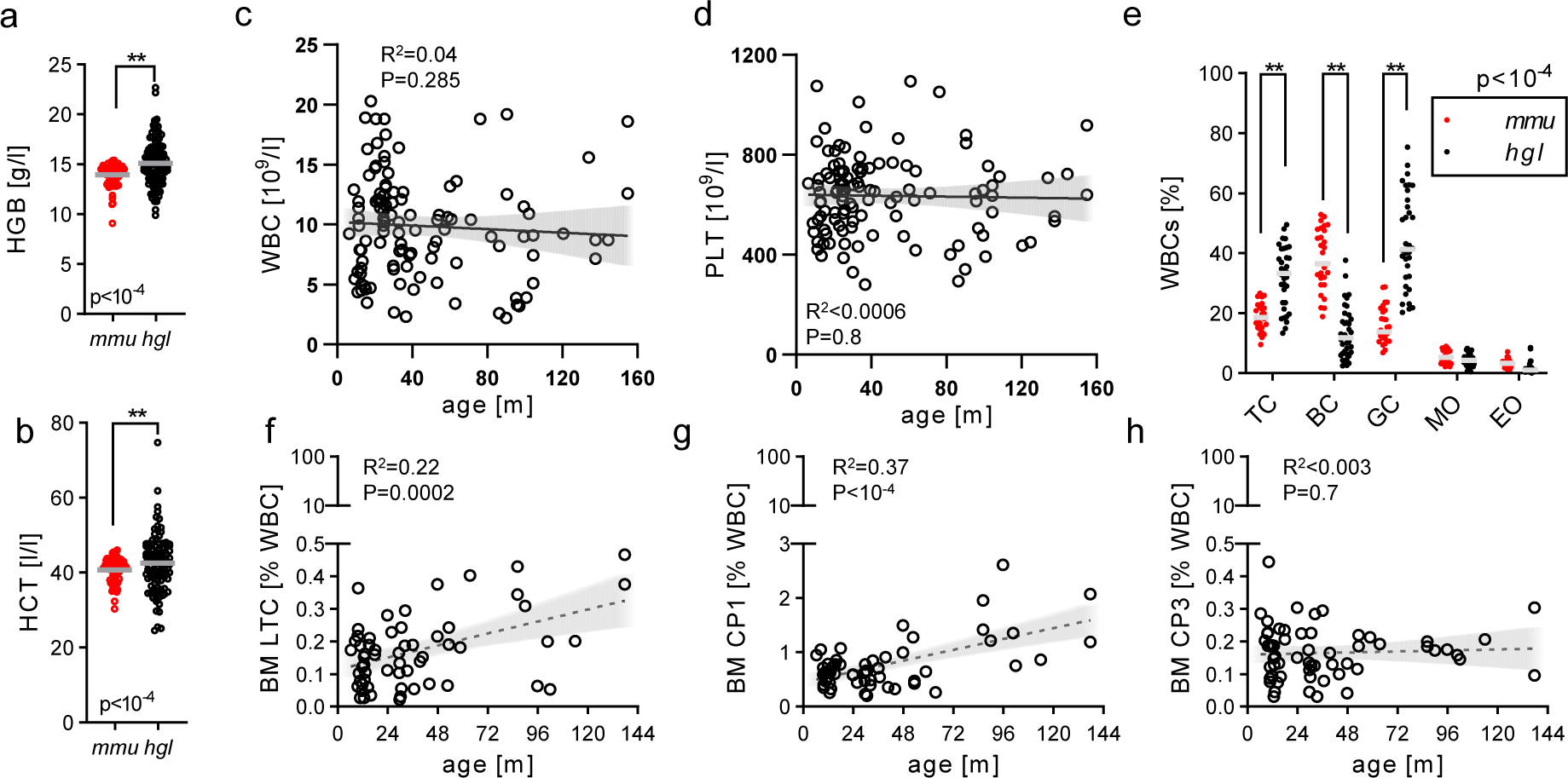
Youthful blood cell composition into midlife in naked mole-rats. **a**, Hemoglobin concentration and **b**, hematocrit levels between mouse [*mmu*] and naked mole-rat [*hgl*] blood. p-values were determined by unpaired Welch’s t-test; n(*mmu*)=83, n(*hgl*)=104. **c**, Volumetric white blood cell (WBC) and **d**, platelet (PLT) numbers across animal age. R^2^, coefficient of determination; p-values were determined by conventional linear regression fitting both slope and intercept; n=112. **e**, FACS WBC frequencies; p-value determined by Sidak’s Two- way ANOVA comparing mouse mmu (n=29) vs hgl (n=34). BM cell frequencies of **f**, LTC/CP2, **g**, CP1 or **h**, CP3 across age; n=60, linear regression with 95% CI as trend line; p<0.05, significance.

In summary, we established a cross-reactive FACS antibody panel to purify HSPC and mature blood cell populations and mapped the major hematopoietic lineages in NMRs.

### The naked mole-rat spleen is the major site of erythropoiesis

One of the first milestones towards prospective isolation of HSCs was the early notion that the cell fraction with hematopoietic regenerative potential was nearly devoid of markers of mature blood cell types (*20*). We thus designed an NMR lineage depletion cocktail (LIN) from the validated cross-reactive antibodies consisting of CD11b/CD18/CD90/CD125, which purified HSPCs as demonstrated by significantly higher colony formation than LIN^+^ or total BM (Fig. 2a-c, Fig. S4c). The LIN^+^ fraction strongly enriched Thy1.1^hi^ GCs, and to a lesser extent TCs and BCs according to their Thy1.1 label intensity in PB populations (Fig. 1b). CD11b and CD18 both form the integrin Mac-1, marking myeloid or NKC commitment. Moreover, we found that the anti-rat Thy1.1 MoAb labels additional cells not commonly stained with anti-human CD90 MoAb in BM, most likely due to different epitopes, each with a proteoform-specific label for NMR THY1 (Fig. S4d).

Scatter backgating demonstrated that CD90^hi^/Thy1.1^hi^ cells were neutrophils, while dim CD90^lo^/Thy1.1^int^ cells had lymphoid scatter properties. We therefore used CD90 antigen to deplete committed cells from the Thy1.1 label. CD125, the IL-5 receptor alpha subunit, is primarily expressed on eosinophils and activated BCs (*21*). Indeed, BCs were the sole fraction positive for CD125 in PB, while TCs, EOs and GCs were gradually labelled by CD90 (Fig. 1b, Fig. S4e). Most cells of the LIN^−^ fraction were Thy1.1^−^/CD34^−^ (CP7; candidate population), resembling committed cells not covered by the NMR LIN cocktail (Fig. 2c). Surprisingly, both LIN^+^ and LIN^−^ contain a Thy1.1^int^/CD34^hi^ population, which we termed CP1 and CP2, respectively. CP3 is LIN^−^ /Thy1.1^lo^/CD34^hi^, CP4-5 are LIN^−^/Thy1.1^−^/CD34^hi^ and LIN^−^/Thy1.1^−^/CD34^lo^, respectively, while LIN^−^/Thy1.1^lo^/CD34^−^ is CP6. Checking which LIN factor is differentially expressed on Thy1.1^int^/CD34^hi^ cells we found CD11b and CD90 absent in CP7 and CP2 but present in CP1 and in most viable Thy1.1^int^/CD34^hi^ cells, all four being negative for CD125 (Fig. S4f). We subset CP1 and distinct cell populations of the LIN^−^ fraction and found each of CP1, CP2 and CP3 were <1% of total BM leukocytes (Fig. 2d), the frequency of the mouse LIN^−^/Sca-1^+^/Kit^+^ (LSK) hematopoietic stem and progenitor cell compartment (Fig. S8c) (*22*).

In most mammals the spleen primarily acts to recycle aged erythrocytes (*23*). However, species-specific adaptations have been found, such as the murine spleen acting as a reservoir of MOs or the equine spleen as a storage of up to 30% RBCs (*24, 25*). We observed a drastic difference in the LIN staining pattern as compared to BM with a strongly expanded LIN^dim^ population corresponding to elevated Thy1.1^−^/CD34^−/lo/hi^ cells in NMR spleens (Fig. S4g). The frequencies of CP4 (2.3-fold), CP5 (3.5-fold) and CP6 were significantly increased in spleens relative to BM (Fig. 2e-f). Likewise, we found increased RBC content in splenic vs marrow organ sections in NMRs but not in mice (Fig. S4a). Reanalysis of scRNA-Seq datasets (*26*) confirmed the progenitors and differentiated cells of the erythroid lineage in NMR spleens, which were absent in mice (Fig. S5, Table S3). Moreover, Benzidine-stained cytospins of whole spleens revealed significantly more RBCs in 3 year-old NMRs than 3 month-old mice (Fig. 2g-h). We further detected Benzidine^+^ nucleated erythroid precursors in NMR but not mouse spleens. Strikingly, in adult mice where normal erythropoiesis is known to occur in the BM, the number of nucleated erythroid progenitors diminished 19-fold from BM to Spleen, in contrast to a 2- fold increase from BM to Spleen in NMRs (Fig. 2h), pointing towards shared splenic and medullary erythropoiesis. To link elevated nucleated erythroid progenitor levels with expansion of the Thy1.1^−^/CD34^lo/hi^ compartment we sorted CP3 and CP4 cells from spleens for Benzidine staining (Fig. 2i). This clearly demonstrated an increase of nucleated erythroid progenitors along with a decline of CD34 expression from CP4 to CP5. Two weeks post-natal is the latest time point in ontogeny where active erythropoiesis takes place in the spleen (*27*), thus continuous utilization of splenic erythropoiesis throughout life can be considered a neotenic trait in NMRs. Consistently, Benzidine- stained colony assays showed an increase in the proportion of hemoglobin-containing colonies from CP1 (0.08) over CP2 (0.26) and CP3 (0.56) to CP4/5 (0.94/0.79; Fig. 2j).

Notably CP1 colonies featured fewer mixed Benzidine^+/–^ colonies than CP2, pointing towards lymphomyeloid lineage restriction of CP1 cells. We thus defined erythroid commitment in the LIN^−^ compartment by a gradual loss of Thy1.1, directly followed by successive downregulation of CD34.

The complete blood counts of NMRs showed higher hematocrit and RBC hemoglobin than mice (Fig. 3a-b). Total RBC numbers were lower than in mice and did not change increase in WBC with age in an NMR cohort spanning 12 years of age (Fig. 3c). Likewise, blood platelet levels increased in mice, but did not increase and were ∼2-fold lower in NMRs (Fig. 3d, Fig. S4k). Hemanalyzer differential platelet counts between the two species were corroborated with RBC:PLT ratios obtained from Wright-Giemsa stained blood smears (Fig. S4l). Imaging of longitudinal femur sections showed fewer erythropoietic islets and Megakaryocytes (MKs) for NMR long bones as compared to mice (Fig. S4a-b).

Using the FACS gating from Fig. 1b we compared the major blood cell types to mice and found dramatically increased granulocytes and reduced BCs in NMRs (Fig. 3e), confirming the higher myeloid:lymphoid ratio (*26*). FACS-based blood cell quantifications were fortified by hemanalyzer measurments (Fig. S4m).

Shared splenic and medullary erythropoiesis may have evolved in NMRs as an adaptation to life in hypoxic conditions (*28*). Additionally, it also provides an alternative functional HSC niche throughout life, which may benefit longevity by sustaining youthful RBC production and preventing age-associated anemia. Moreover, NMRs did not display age-associated increase of blood leukocytes and platelets, pointing towards reduced chronic inflammation and delay of age-associated thrombosis.

### LTCs (LIN^−^/Thy1.1^int^/CD34^hi^) are the main source of naked mole-rat HSCs

We next performed population RNA-Sequencing of sorted CP1-4 fractions to annotate their developmental status. Unsupervised clustering by t-distributed stochastic neighborhood embedding (t-SNE) separated transcriptomes in accordance with their immunophenotype (Fig. 4a). Due to the transition from LIN^−^ to LIN^+^ between CP2 and CP1 we checked which genes were successively downregulated during transition from LTCs (CP2) to CP1 and CP3/4 (Fig. 4b). We retrieved 116 genes showing this expression pattern, of which 40 are found in the LTC RNA-Seq signature (Table S4). A key finding was high expression of ID2, which blocks BC differentiation, can enhance erythropoiesis and expands HSCs (*29, 30*). LTCs also showed high expression of CD81, a tetraspanin which has been shown to maintain self-renewal in HSCs (*31*). Notably TM4SF1, the top marker of NMR HSPCs from the scRNA-Seq atlas (Fig. 1h), and the pluripotency marker EPCAM which facilitates reprogramming (*32*), are most abundant in LTCs.

**Fig. 4.**
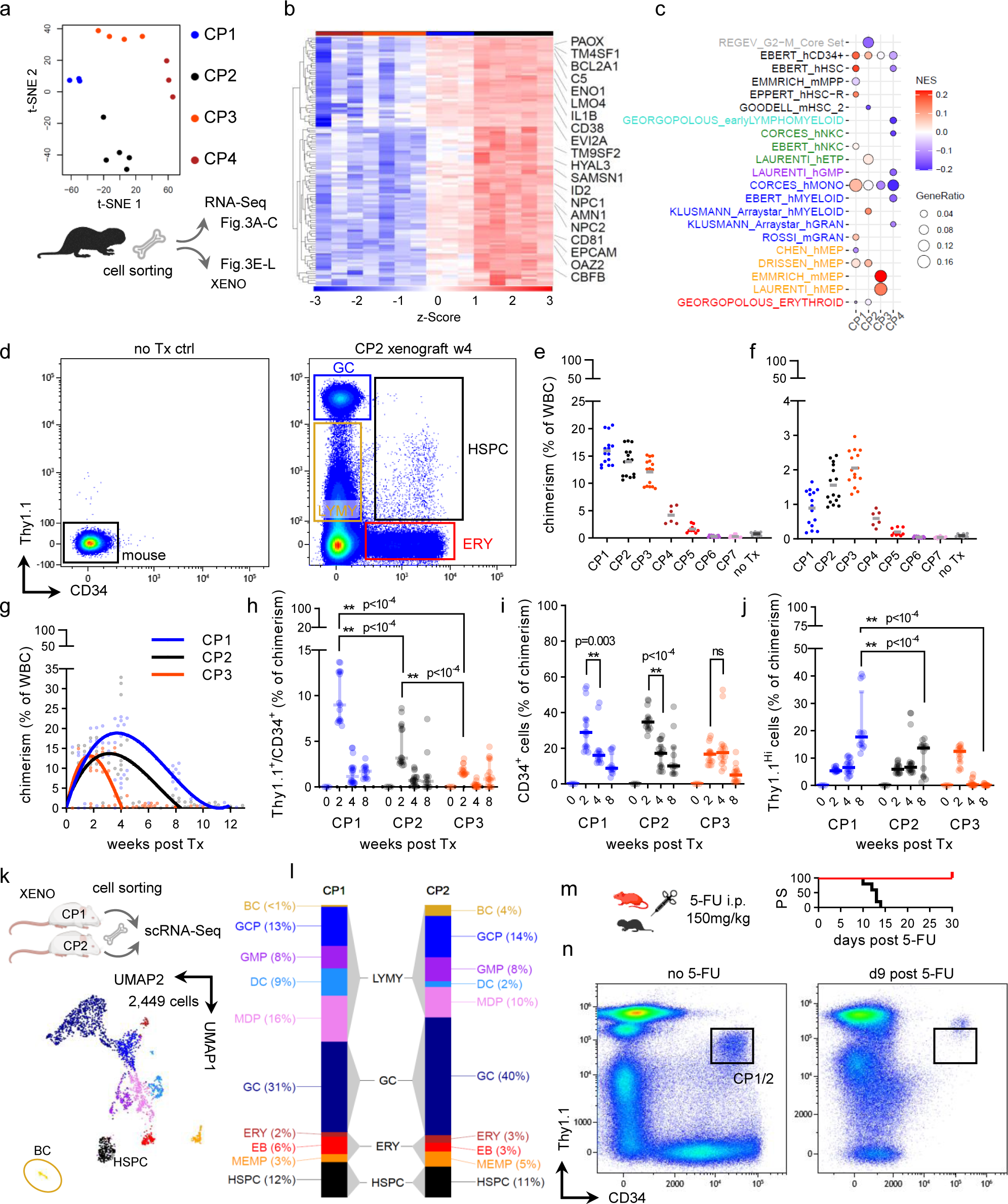
LTCs define the primitive HSPC compartment. **a**, BM from 1-3 year old naked mole-rats sorted into indicated cell populations for RNA- Sequencing or xenotransplantations, color legend applied throughout the Figure. Unsupervised t- SNE clustering, effectively separating each CP group; vst-transformed counts as input. **b**, 116 gradually downregulated genes from CP2 to CP1, displayed are 20 genes with known roles in hematopoiesis. **c**, GSEA of sorted BM fractions displaying top 10 q-value terms from a geneset collection of human and mouse HSPCs. NES, normalized enrichment score; GeneRatio, (signature ∩ term) / (signature ∩ all terms). **d**, Gating strategy to quantify total engraftment; untransplanted recipient [left] vs CP2 xenograft 4 weeks post Tx [right]; HSPC, Thy1.1^int^/CD34^+^ stem and progenitors; ERY, CD34^+^ erythroid cells; GC, Thy1.1^hi^ granulocytes; LYMY, Thy1.1^lo/int^/CD34^−^ lymphomyeloid cells. Recipient chimerism 2 weeks post Tx in **e**, BM or **f**, spleen; total recipients from 3 donors for each CP graft. **g**, BM chimerism over time; recipients from 5 donors for weeks 4, 8 and 12. P-value determined by Fisher’s Two-way ANOVA; curve- fitting by cubic polynomial. Kinetics of engraftment proportions for **h**, HSPC, **i**, ERY and **j**, GC; p-value determined by Tukey’s Two-way ANOVA. **k**, ScRNA-Seq of week 4 BM CP1 and CP2 xenografts (n=1); HSPC and BC clusters are outlined in CP1/2-integrated dataset filtered for naked mole-rat cells. **l**, Quantification of GSEA-annotated cell types between grafts. **m**, 5-FU administration into 6 month-old mice (n=4, bodyweight 25±2g) or 2-3 year-old naked mole-rats (n=5, bodyweight 29±4g); i.p., intraperitoneal. **n**, Naked mole-rat BM untreated or treated with 5- FU (n=5), no LIN gating.

Next we derived differentially expressed genes specific to each CP1-4 (Table S4). Strikingly, the strongest GSEA association for CP3 were MEPs, while CP4 is negatively expression of erythroid marker genes GATA1, EPOR, TFRs, KEL and FECH in CP3/4 (Fig. 4c, Fig. S6a). CP1 was enriched with the most HSPC-associated genesets (Table S4). The human CD34^+^ signature displayed an enrichment gradient from CP1 to CP4, revealing a Mac-1^+^ primitive progenitor fraction with a stemness expression profile in adult NMRs.

The capacity to give rise to several distinct lineages via differentiation, referred to as multipotency, can be assayed through quantitation of progenitor frequencies during colony formation (*33*). We observed that NMR HSPCs grew best at 32°C in methylcellulose supplemented with human cytokines (Fig. S6b). Scoring of colony-forming unit (CFU) types was validated by cytochemistry of single colonies (Fig. S6c-d). Of all NMR BM populations only CP6 and CP7 did not grow in methylcellulose assays (Fig. S6e).

Furthermore, the proportion of erythroid over total colonies declined from CP4/5 (0.94/0.81) to CP3 (0.53), CP2 (0.29) and was lowest in CP1 (0.2). Myeloid output was not significantly different between CP1 and CP2 but decreased in CP3. Serial replating yielded 1.5-fold more total colonies for CP2 compared to CP1, although no colony type frequency was significantly altered between these two, as seen for original platings (Fig. S6f). Multipotency can further be assessed by transplantations into preconditioned immunodeficient hosts, through which high levels of sustained primary engraftments could be obtained in a variety of humanized mouse models (*34*). Given the successful *in vitro* growth of NMR HSPCs with human cytokines we reasoned that the NSGS mouse model with constitutive production of human IL-3, M-CSF and SCF would render optimal support to NMR xenografts (*35*). We indeed observed robust engraftment rates for CP2 at 4 weeks post transplantation in recipient BM as compared to untransplanted mice (Fig. 4d). Xenografts recapitulated the FACS staining pattern of naked mole-rat BM origin and could be separated from host cells, which are not labelled by validated NMR Thy1.1 and CD34 moAbs (Fig. S6g-j). At week 2 host BM chimerism resembled colony yields from methylcellulose assays with CP6/7 engraftments below background, supporting the notion that the NMR HSPC compartment is CD34^+^ as in humans (Fig. 4e). All other populations produced clearly detectable engraftment in NSGS mice ranging from 1.6% (CP5) over 4.2% (CP4) and 12.1% (CP3) to 14% (CP2) and 16% (CP1). Repopulation of host spleens was markedly reduced for all engrafted groups; strikingly CP3 was functionally classified as the most primitive committed erythroid progenitor and enriched in host spleens with higher engraftment than CP2 (Fig. 4f). Though FACS analyses verified xenograft cells in shown). BM engraftment at week 4 for CP3 (1%) depleted earlier than for CP1 (12.8%, p<10^-4^) and CP2 (11.7%, p<10^-4^) (Fig. 4g). The early loss of erythroid-primed CP3 is consistent with higher residual chimerism at week 8 for myeloid-primed CP1 compared to CP2 (5.3% vs 1.3%, p=0.04). Unexpectedly none of the three most primitive stem and progenitors or whole marrow (WBM) showed sustained BM engraftment past 12 weeks (Fig. S6k), a fact we primarily attribute to the difference in body temperature between NMRs (thermoneutral at ∼32°C) and humans or mice, leading to niche stress on the graft and its depletion.

Next we quantified lineage commitment over time by selecting the Thy1.1^+^/CD34^+^ compartment of xenografts (HSPC, Fig. 4d). Although this rapidly depleted for all cell types at week 4, the initial replicative burst was greater in CP1 compared to CP2 (Fig. 4h), suggesting that CP1 cells are activated to a greater extent by the inflammatory host environment that ultimately exhausts engraftment. Xenograft CD34^+^ cells (ERY, Fig. 4d) resembling the erythroid lineage decline towards week 4 for CP1 and CP2, whereas CP3 CD34^+^ output remained similar (Fig. 4i). Conversely, we used Thy1.1^hi^ cells as myeloid output (GC, Fig. 4d), which revealed most efficient myelopoiesis at week 8 in CP1 compared to CP2 and CP3 (Fig. 4j). B-lymphopoiesis in NMR BM is conserved (Fig. 1k, Fig. S2e, 3e), and since blood BCs are labelled by Thy1.1^lo^/CD11b^−^, we reasoned that CP6 cells would contain marrow and spleen BCs, albeit with less purity. The xenograft lymphomyeloid population (LYMY, Fig. 4d) significantly dropped in CP3 cells at weeks 4 and 8 but is more efficiently sustained in CP1 and CP2 with higher myeloid potential (Fig. S6l). Since the heterogeneity of this FACS fraction does not provide evidence over definitive B-lymphoid commitment in xenografts, we performed scRNA-Seq from week 4 CP1 and CP2 grafts (Fig. 4k). Integrated analysis on both grafts identified a rare BC population amongst erythroid cells, HSPCs and 75% myelocytes (Fig. 4l, Fig. S6m, Table S5). Considering the exhaustive effect of the host BM niche a myeloid bias under stress hematopoiesis and inflammatory conditions is expected. CP2 clearly produced almost all BCs (6.5-fold to 4% total xenograft compared to 0.6% CP1), suggesting that lymphoid commitment within the primitive HSPC compartment is lost upon Mac-1/CD90-antigen expression at the onset of myelopoiesis in CP1. Concordantly, the CITE counts for CD11b corresponded with LIN sorting between CP1/2 (Fig. S6n).

5-Fluorouracil (5-FU) eliminates cycling hematopoietic cells and activates the dormant myeloablation in mice. In same-sized NMRs however this dose led to complete mortality before day 15 post administration (Fig. 4m). At day 9 when BM is almost completely reconstituted in mice (*37*), the entire CD34^+^ compartment in NMRs was lost leaving an aberrant LIN^+^/Thy1.1^hi^/CD34^hi^ marrow GCP fraction (Fig. 4n). In terminal anemic animals erythroid Thy1.1^−/lo^/CD34^+^ fractions were not regenerated (Fig. S7a), strongly supporting that LTCs, which are not restored upon ablation, contain *bona fide* HSCs.

Given a stronger Rhodamine 123 (Rho) efflux, which functionally enriches human and mouse HSCs (*38, 39*), in LTCs than in LSKs (Fig. S7b), the sensitivity to 5-FU is not caused by impaired drug transporter systems.

Altogether CP1 cells resemble CMPs, albeit with expression of Mac-1, CD90-antigen and a human-like HSPC signature. The lack of lymphoid development in xenografts points towards a primitive myeloid progenitor with severely decreased capacity of differentiating towards the erythroid lineage. Furthermore, we functionally defined the most primitive HSPC compartment as CP2 LTCs, exhibiting the highest degree of quiescence and multipotency. Our FACS panel effectively subsets the primitive HSPC compartment in NMRs, wherein diminished Thy1.1 expression of CD34^+^ cells correlated with erythroid fate decision along the LTC-CP3/4 axis, while rising CD11b levels corresponded to myelopoiesis through LTC-CP1.

### Expansion of marrow granulopoiesis and the erythroid lineage at the expense of B-lymphopoiesis

Next we ran CITE-Seq on WBM from two 3 month-old and two 12 month-old mice (*mmu*) against WBM from two 3 year-old and two 11 year-old naked mole-rats (*hgl*; Fig. 5a, Fig. S7c). We used canonical correlation analysis to integrate the four marrow libraries separately for each species (*40*). Louvain clustering found 15 communities from a total of 19298 mouse marrow cells; conversely 14 communities in 21678 NMR marrow cells were detected (Fig. 5b), cluster annotation based on GSEA (Table S5). In mice cell types were strongly aligned with the CITE signals (Fig. S4d), e.g. a rare HSPC population of <1% total BM expressed ANGPT1, GATA2 and HOXA9 and had CITE-LIN^−^/Kit^+^/Sca-1^−/+^, co- clustering myeloid progenitors (LKs) and LSKs as the murine HSPC compartment.

**Fig. 5.**
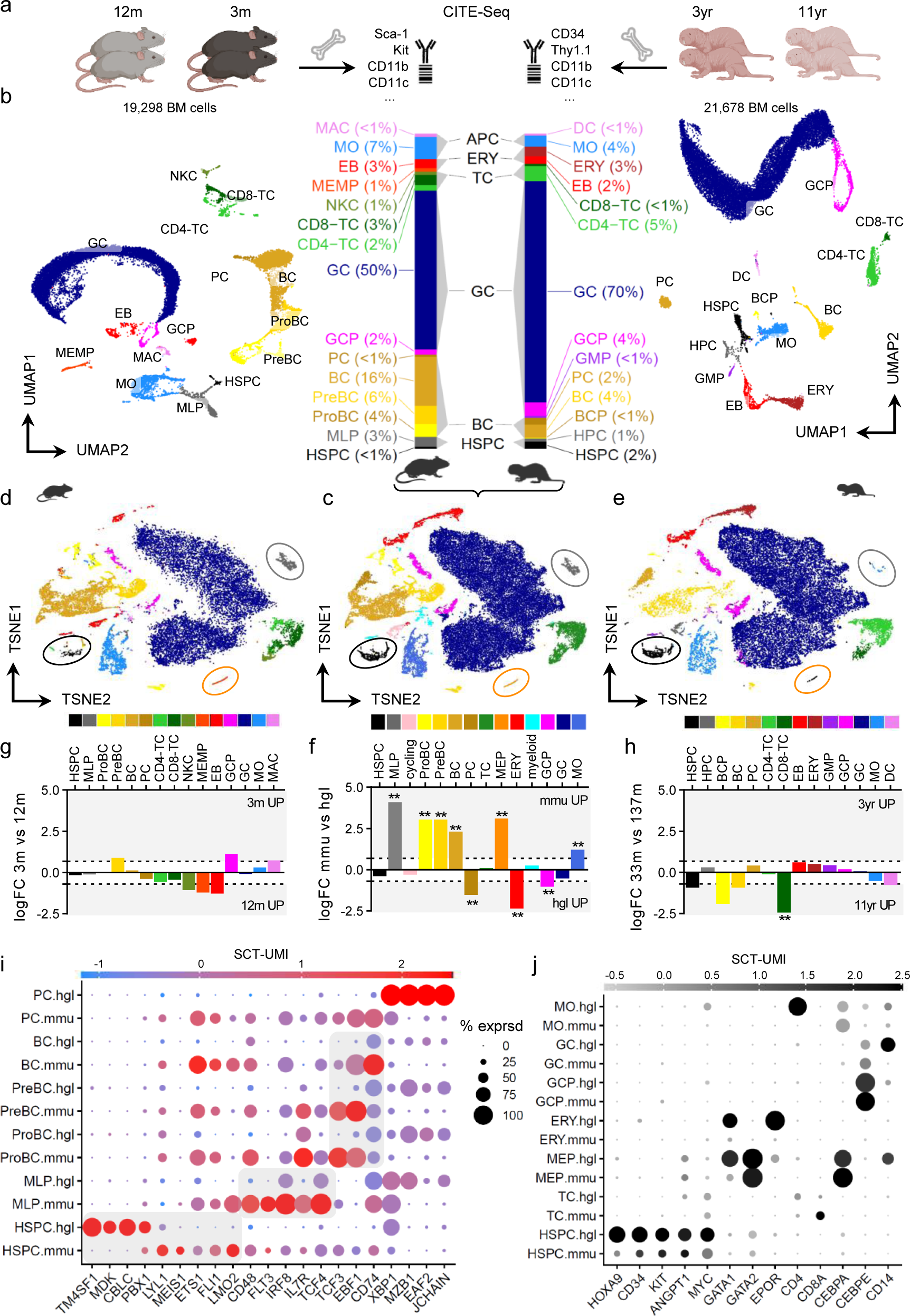
BM CITE-Seq across species. **a**, Unfractionated BM of perfused mice (n=4) or naked mole-rats (n=4) were subjected to CITE- Seq. *mmu*CITE-moAbs: Sca-1, Kit, Cd11b, Cd11c, Nk-1.1, Cd4, Cd8a, Cd3e, Cd19, Cd25, Cd44, Gr-1, Ter119; *hgl*: Thy1.1, CD34, Cd11b, Cd11c, Nk-1.1, Gr-1, CD90. **b**, UMAP of the CCA-integrated mouse [left] or naked mole-rat [right] dataset with fGSEA-annotated cell types. MEMP, megakaryocytic erythroid mast cell progenitor; GCP, granulocytic precursor; HPC, hematopoietic progenitor cell. Bar chart [center] displaying average cluster frequencies across species; APC, antigen-presenting cell. **c**, T-SNE of SCTransform-integrated *mmu* and *hgl* BM, colorbar legend for species-integrated clusters below. Encircled coordinates for HSPC, LMPP and MEP clusters; cycling, co-clustered based on active cell cycle gene expression. T-SNE from species-integration for **d**, mouse or **e**, naked mole-rat partition; Cluster annotation and coloring from the single-species analysis in b. **f**, Differential cell type abundance across species; dotted lines, 2-fold change. MLP, p<10^-10^; ProBC, PreBC, MEP, p<10^-7^; BC, ERY, p<10^-5^; PC, p=0.0017; MO, p=0.0088; GCP, p=0.027. Differential cell type abundance across age for **g**, mouse or **h**, naked mole-rat single-species analysis. *hgl* CD8-TC, p=0.002; *hgl* BCP, p=0.013. Dotted lines, 2-fold-change. Cross-species expression of selected **i**, differentially regulated or **j**, conserved markers between cell types. SCT-UMI, sctransform-scaled UMI counts; % exprsd, percentage of cells/cluster with UMI ≥ 1.

Likewise, NMR HSPCs overexpressed TM4SF1, GATA2 and HOXA9, and were CD11b^−/lo^/CD34^+^/Thy1.1^−/lo/int^, suggesting LTCs, CP1 and CP3 collectively clustered as HSPC. Next we quantified cell types across species and grouped them into major branches (Fig. 5b). The most abundant fractions were GCs and granulocytic precursors (GCP), expressing conserved cell type markers, which account for ∼50% WBM in mice and ∼75% in NMRs. By contrast, the BC compartment with >26% in mice is reduced 4-fold to <7% in NMRs. As reported previously for spleen(*26*), there are no NKCs detected in NMR BM. Surprisingly, although the TC partition has the same frequency in both species, mice show a 2:3 ratio of CD4- vs CD8-TCs, NMRs a ratio of 5:1. In fact, by absolute copy number quantitation of sorted PB-TCs we determined a CD4:CD8 mRNA ratio of 1:3.5 in mice, which was inverted to 2.5:1 in NMRs (Fig. S7e). This pattern was confirmed in whole LNs with mature peripheral TCs as the primary source of CD4/CD8 expression, showing a 1:2.2 ratio in mice and 5.8:1 in NMRs (Fig. S7f). Hereby NMRs resemble humans rather than mice. The CD4:CD8 ratio is used to discriminate the risk of disease progression in HIV/AIDS and decreases with age in patients (*41*), a low CD4:CD8 ratio indicates immunosenescence and is associated with wide-ranging pathology (*42*). The unusually high CD4:CD8 ratio in NMRs suggests a reduced dependence on cell-mediated immunity.

Assuming a linear relationship between age groups across species, we compared differences in gene expression and cell type abundances of 12 month-old versus 3 month-old mouse BM with 11 year-old versus 3 year-old NMR BM. In mice no cluster abundance was significantly altered across age groups (Fig. 5g), however GSEA on 760 differentially expressed genes across all clusters between age groups revealed 3 upregulated terms related to proliferation and growth in older mice (Fig. S7g). Only CD8- TCs were significantly elevated in older NMRs to 1% of total BM cells (11-fold, q<0.033; Fig. 5h), which most likely reflected memory cell acquisition over age. There were no significantly associated pathways from the 978 differentially expressed genes. While we did not expect to find strong age-associated differences in gene expression between 3 month-old and 12 month-old mice, long-lived NMRs retained a youthful BM composition at least for the first decade of their lifespan.

Next we integrated both species data using SCTransform (*43*), and overlaid species annotations on integrated clusters (Fig. 5c-f). We found 91.4% of mouse HSPCs and 85.8% of NMR HSPCs mapping to the same integrated HSPC cluster, which commonly expressed stem cell markers such as HOXA9, KIT and ANGPT1 (Fig. 5j). Though this could lead to conclude a higher HSPC abundance in NMRs (0.54% *mmu* vs 2% *hgl*; Fig. primitive HSPC compartment of both species, supported by transcriptional signatures of human HSC and CD34^+^ cells associated with CP1 (Fig. 4c). Conversely, CITE-CD11b^+^ HSPC cluster cells increased in older animals (Fig. S7h), and the HSPC cluster of older NMRs was expanded by ∼2-fold, albeit not significant (Fig. 5h). Strikingly, both CP1 and CP2/LTC BM frequencies significantly increased with age, whereas erythroid progenitors remained constant (Fig. 3f-h, Fig. S4n). Therefore an intrinsic myeloid differentiation bias progressing with age is inherent to NMR HSPCs, akin to human HSCs (*44*).

Surprisingly, there were no NMR counterparts co-clustering with murine multipotent lymphoid progenitors (MLPs) within the integrated MLP community, thus mouse BM strongly enriched for MLPs (59-fold, q<10^-9^; Fig. 5f). Mouse MLPs had lower levels of stem cell factors Lmo2 and Pbx1 than HSPCs, instead shared markers upregulated throughout the B-lineage such as Ets1, Fli1, Cd48, Il7r, Tcf3, and showed specific overexpression of Tcf4, Irf8 and Flt3 (Fig. 5i).

NMR cells mapping to the integrated MLP population were annotated as MOs in the single-species clustering (Fig. 5e) and overexpressed IRF8 and TCF4 but not IL7R or LMO2. As the earliest HSPC-to-B-transition intermediate MLPs were accumulated along with ProBCs, PreBCs (both 21-fold, q<10^-6^) and BCs (10-fold, q<10^-5^) in mouse BM (Fig. 5f). Evidently the significant B-lineage reduction in NMRs manifests from primitive progenitors to mature BCs. However, in NMR BM a rare population with high expression of JCHAIN, MZB1, XBP and EAF2 likely comprised germinal center B or plasma cells (PC; Fig. 1h, Fig. S5, 7d). Intriguingly, there are more PCs in NMR BM than in mice (Fig. 5f). The higher CD4-TC abundance combined with a compressed BC compartment leads to a higher CD4:APC (antigen presenting cell) ratio and could more efficiently activate BCs, relative to their total frequency, resulting in more plasmablastoid differentiation. On the other hand, increasing the amount of the terminal effector cell, e.g. through lower cell turnover, could have evolved to compensate a less abundant BC compartment.

Mouse megakaryocytic/erythroid/mast cell progenitors (MEMP) form a distinct cluster in mouse whole marrow, which is reformed upon integration of both species (MEP), of which the NMR fraction derived from HSPCs (Fig. 5b-e). We observed splenic erythropoiesis as the primary route to NMR RBC production, however the increase of erythroid cells in BM (11-fold, q<10^-5^) suggested a higher prevalence of erythroid commitment (Fig. 5f). Surprisingly mouse BM contained 22-fold more MEPs (q<10^-6^), supported by GATA1/GATA2 co-expression common in MEPs of both species (Fig. 5j).

MEPs, whereas integrated naked mole-rat MEPs expressed GATA2^+^/GATA1^hi^ (Fig. 5j). This is further in line with reduced BM MKs and PB platelets in naked mole-rats.

In summary we have shown that naked mole-rat BM maintains a myeloid bias towards granulopoiesis, accompanied by reduced B-lineage commitment. A stem cell state was portrayed by pervasive expression of TM4SF1, the top HSPC marker by fold-change, highest expression in LTC and most specific for NMR HSPCs across species.

Erythropoiesis is favored over megakaryopoiesis in NMR marrow, contributing to maintenance of low platelet levels in PB.

### Naked mole-rat HSPCs display low metabolism and slow cell cycling

Since CITE-Seq driven cell type annotations matched HSPC FACS populations, we examined sorted population level transcriptomes of corresponding developmental stages across species (Fig. 6a, Fig. S8a-e). A comprehensive collection of distinct human and murine HSPC stages was retrieved from GEO and integrated with bulk RNA-Seq data from human and NMR. The dataset of 9422 orthologs across 218 transcriptomes was segregated into 3 groups, primitive (LT-HSC, MPP), lymphomyeloid and erythroid progenitor. Using GSEA with MSigDB hallmark genesets we found that mouse cells through all stages were enriched in mitotic and pro-proliferative pathways (Fig. 6a).

**Fig. 6.**
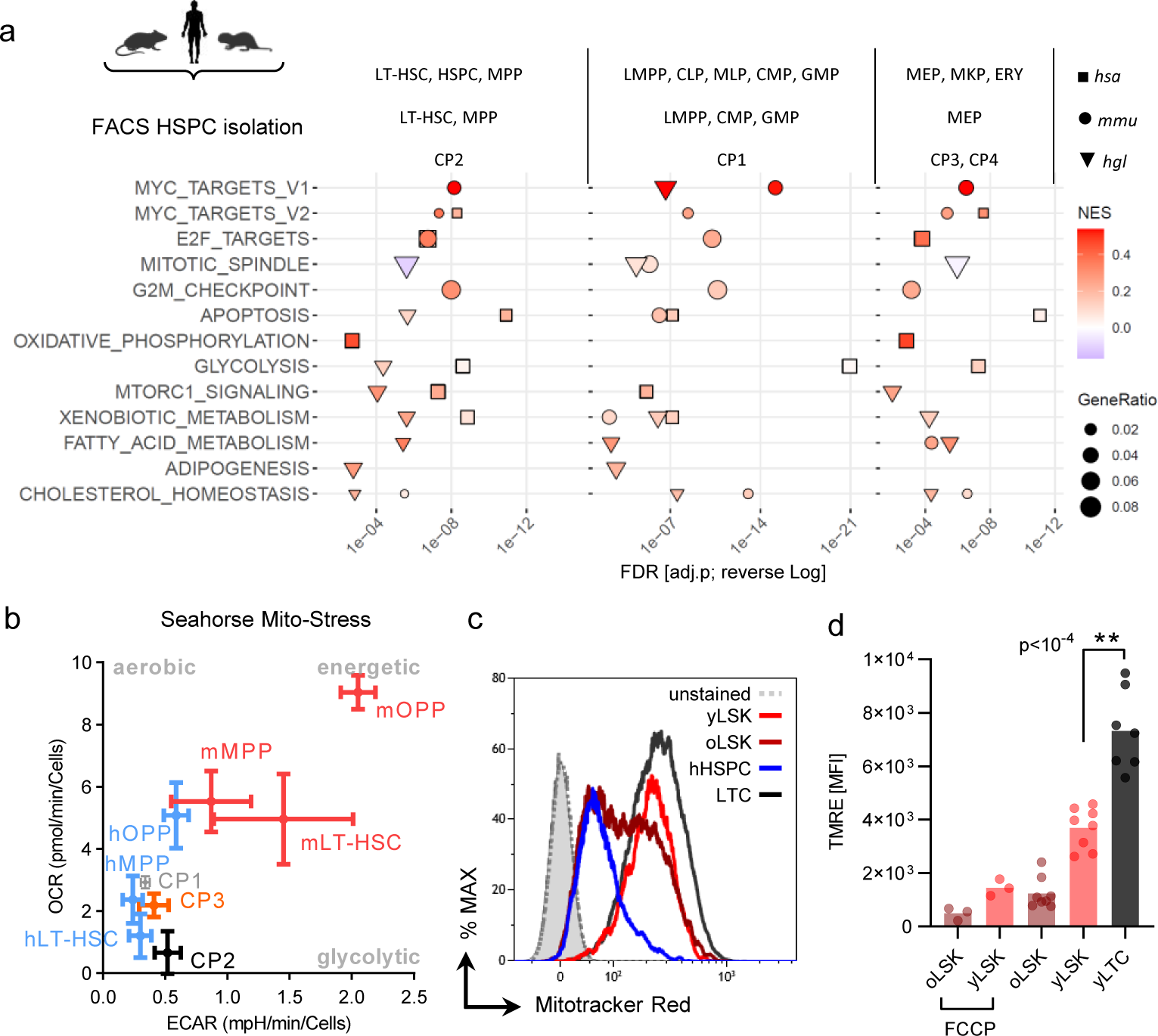
Low metabolism of naked mole-rat HSPCs. **a**, Cross-species integration of bulk RNA-Seq datasets. Naked mole-rat BM populations CP1-4 were matched to human and murine counterparts for GSEA with MSigDB hallmark genesets. Primitive stem and progenitor [left], *hsa*: LT-HSC, LIN^−^/CD34^+^/CD38^lo^/CD90^+^/CD45RA^−^ (n=7); HSPC, LIN^−^/CD34^+^/CD38^lo^ (n=11); MPP, LIN^−^/CD34^+^/CD38^lo^/CD90^−^/CD45RA^−^ (n=4); *mmu*: LT-HSC, LIN^−^/Sca-1^+^/Kit^+^/CD150^+^/CD48^−^ (n=25); MPP, LIN^−^/Sca-1^+^/Kit^+^/CD150^−^/CD48^+^ (n=6); *hgl*: CP2, LIN^−^/Thy1.1^int^/CD34^+^ (n=5). Lymphomyeloid [center], *hsa*: LMPP, LIN^−^ /CD34^+^/CD38^lo^/CD123^lo^/CD45RA^+^ (n=9); CLP, LIN^−^/CD34^+^/CD38^hi^/CD10^+^/CD45RA^+^ (n=6); MLP, LIN^−^/CD34^+^/CD38^lo^/CD90^−^/CD45RA^+^/CD71^−^ (n=4); CMP, LIN^−^ /CD34^+^/CD38^hi^/CD123^lo^/CD45RA^−^ (n=26); GMP, LIN^−^/CD34^+^/CD38^lo^/CD123^lo^/CD45RA^+^ (n=18); *mmu*: LMPP, LIN^−^/Sca-1^+^/Kit^+^/Flt3^hi^ (n=5); CMP, LIN^−^/Sca-1^−^/Kit^+^/CD16/32^lo^/CD34^+^ (n=8); GMP, LIN^−^/Sca-1^−^/Kit^+^/CD16/32^hi^/CD34^+^ (n=16); *hgl*: CP1, LIN^+^/Thy1.1^int^/CD34^+^ (n=5). Erythroid [right], *hsa*: MEP, LIN^−^/CD34^+^/CD38^hi^/CD123^−^/CD45RA^−^ (n=23); MKP, LIN^−^ /CD41a^+^/CD42b^+^ (n=3); ERY, LIN^−^/CD34^lo/–^/CD36^+^/GYPA^+^/CD71^+^ (n=12); *mmu*: MEP, LIN^−^ /Sca-1^−^/Kit^+^/CD16/32^−^/CD34^−^ (n=15); *hgl*: CP3, LIN^−^/Thy1.1^lo^/CD34^+^ (n=5); CP4, LIN^−^/Thy1.1^−^/CD34^+^ (n=5). Within each group, cell types were pooled for each species. Shown are pathways related to proliferation and metabolism, full results see Table S6. NES, normalized enrichment score; GeneRatio, (signature ∩ term) / (signature ∩ all terms); FDR, false discovery rate. **b**, Seahorse assay with cell types sorted according to (A); hOPP, human oligopotent progenitors LIN^−^/CD34^+^/CD38^+^; mOPP, LIN^−^/Sca-1^−^/Kit^+^. ECAR, extracellular acidification rate; OCR, oxygen consumption rate. Human cell types, n=4; mouse, n=3, naked mole-rat, n=3. **c**, Mitotracker Red staining in mouse (n=4), human (n=4) and naked mole-rat (n=5) BM, histogram of merged per-species data. unstained, BM from mouse [solid], human [dotted] or NMR [AUC]; yLSK, LIN^−^/Sca-1^+^/Kit^+^, 3 month old; oLSK, 24 month; yLTC, LIN^−^/Thy1.1^int^/CD34^+^, 3 year old; hHSC, human CD34^+^/CD38^lo^; quantitation see Fig. S8h. **d**, Mean fluorescence intensities of TMRE stainings. p-values were determined by Tukey’s One-way ANOVA; LSK, n=8; LTC, n=7. FCCP, negative control.

Interestingly, functional annotation of scRNA-Seq cluster signatures across species showed mouse HSPCs and several more committed cells enriched in pro-proliferative pathways over their NMR counterparts (Fig. S7i). Human HSPCs scored high for apoptosis, glycolysis and OXPHOS pathways, whereas NMR HSPCs strongly enriched for adipogenesis, cholesterol homeostasis and fatty acid oxidation (FAO) related terms. Indeed a plasma metabolite signature of multiple upregulated lipid sub-classes have been reported earlier (*45*). FAO provides the substrates for OXPHOS, while Aldehyde dehydrogenases (ALDH) neutralize aldehydes arising from processes such as lipid peroxidation. Notably, ALDH staining revealed 2-fold higher levels in LTCs compared to LSKs (Fig. S8f), indicating a countermeasure against elevated FAO activity.

Metabolic paradigms of HSCs are their reliance on glycolysis and low mitochondrial activity (*46, 47*). We measured mitochondrial respiration and glycolysis in sorted HSPCs from 3 species by the Seahorse assay (Fig. 6b, Fig. S8g). We found that human long-term (LT) HSCs and LTCs had the lowest metabolic profile, resembling quiescent cells. Mouse cells showed 2-4-fold higher respiration (OCR) and glycolysis-driven acidification than their human and NMR counterparts, suggesting that quiescent mouse LT-HSCs have a higher basal metabolic rate. The mitochondrial mass of young mouse and NMR HSPCs are similar, whereas human HSPCs have less mitochondria (Fig. 6c, Fig. S8h). The mitochondrial membrane potential (MMP) as a resultant of OXPHOS and FAO is an indicator of mitochondrial activity. We reproduced (*48*) that old LSKs feature an increased fraction of cells with low MMP compared to young LSKs (Fig. S8i). Increased MMP in LTCs compared to LSKs was shown using Tetramethylrhodamine (TMRE) sequestration (Fig. 6d). Superoxide levels in LTCs were lower than those of LSKs (Fig. S8j). In line with seahorse metabolic profiles, intracellular reactive oxygen species (ROS) levels were higher in LSKs than in LTCs and human HSCs (Fig. S8k).

Within NMR HSPCs, a signature of actively cycling cells was enriched for CP1 and depleted for LTC (Fig. 4c). Pyronin Y staining confirmed more G0 LTCs than CP1 (1.5- fold; Fig. 7b). Cell cycle scoring of BM scRNA-Seq clusters revealed 2-fold more mouse HSPCs in S phase compared to NMR HSPCs (Fig. 7a). Conversely, Ki67 staining showed a higher LTC G0 fraction as compared to CP1 (3.4-fold) and mouse HSPCs (Fig. 7b). Next we performed Dual-Pulse labelling (*49*) by successive injection of EdU and BrdU to compare cell cycle kinetics *in vivo* (Fig. 7c). Using the EdU label together with DNA content staining we found a 3-fold increase in S-Phase LSKs over LTCs (p<10^-4^; Fig. 7d, Fig. S9a-b). Committed progenitors of either species did not differ in their cell cycle properties. Combined subsequent use of 2 label incorporations allows quantitation of the cells entering S phase, excluding cells retaining the 1^st^ label to purify cells in early S phase via the 2^nd^ label (*50*). Cells in early S phase incorporate only BrdU (EdU^−^BrdU^+^). Cells in mid/late S phase are at DNA synthesis during both label administrations (EdU^+^BrdU^+^).

**Fig. 7.**
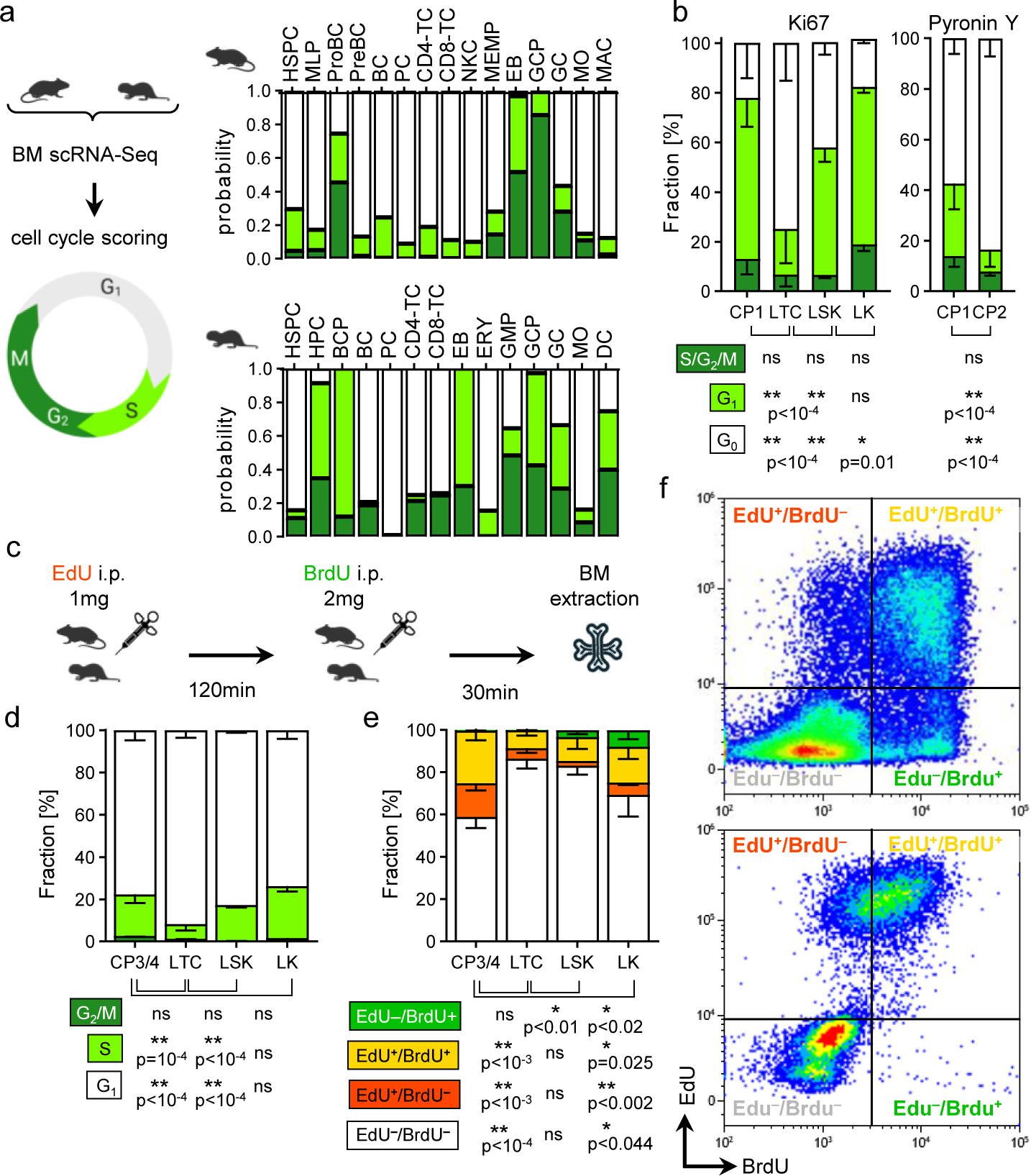
Slow cell growth of naked mole-rat HSPCs. **a**, Cell cycle scoring of mouse [top] and naked mole-rat [bottom] whole BM scRNA-Seq. **b**, Cell cycle staining with Ki67 [left] (CP1/LTC, n=12); young mouse BM (n=4) was used for LK and LSK. p-value determined by Tukey’s Two-way ANOVA. Note that CP1 cells do not differ in any cell cycle stage from LKs. Cell cycle staining with Pyronin Y [right] (N=6); p-value determined by Sidak’s Two-way ANOVA. **c**, Dual-Pulse labelling *in vivo* with 3-4m old mice (n=7) or 2-3yr old naked mole-rats (n=5). EdU, 5-ethynyl-2′-deoxyuridine; BrdU, 5-bromo-2′-deoxyuridine. **d**, Cell cycle analysis of BM populations by EdU vs DNA-content. LK, LIN^−^/Sca-1^−^/Kit^+^; CP3/4, LIN^−^/Thy1.1^lo/–^/CD34^+^; p-values obtained from Tukey’s Two-way ANOVA. **e**, Dual-Pulse analysis of BM populations by EdU vs BrdU. P-values obtained from Tukey’s Two-way ANOVA. **f**, Representative quantitation of merged per-species Dual-Pulse measurements for mouse LK [top] and naked mole-rat CP3/4 [bottom].

Cells post S phase between the two labels incorporated only the first label (EdU^+^BrdU^−^). As expected, the CP3/4 progenitor partition showed markedly more mid/late and post S phase cells than the LTC stem cell compartment (Fig. 7e). Accordingly, the same pattern can be seen for mouse myeloid progenitor LKs versus LSK HSPCs. However, virtually all NMR cells did not show EdU^−^BrdU^+^ early S phase cells (Fig. S9c-d). We thus conclude that the S/G2/M period in NMR cells extends beyond the typical 4h in mice (*51*).

Consequently, even the highly proliferative CP3/4 fraction did not feature early S cells with the 2h between-label interval as compared to LKs (Fig. 7f), thus showing prolonged G1-S progression in NMRs.

Taken together, these data suggest that NMR HSPCs have evolved a mechanism of stem cell homeostasis involving elevated MMP and ALDH activity while maintaining a larger quiescent fraction of their total stem cell pool. OXPHOS through FAO is more energy- efficient and prevents lactate-caused cytoplasmic acidification, which likely contributes to preservation of quiescence and tissue homeostasis during aging.

## Discussion

Naked mole-rats are the longest-lived rodents, and remarkably they remain healthy until the end of their lives and are resistant to age-related diseases including cancer. Adult stem cells are essential for maintenance and repair of tissues, thus NMR stem cell biology is of immediate interest to biomedical research.

Here we present a comprehensive analysis of the blood system in >100 NMRs including functional and molecular characterization of stem and progenitor subtypes and a primary landscape of the hematopoietic hierarchy. Surprisingly, many characteristics of the NMR hematopoietic system showed higher similarity to humans than to mice (Fig. 8). It had been proposed that long-lived NMRs, as well as humans, display neotenic traits compared to their short-lived relatives (*52*). Neoteny describes the preservation of juvenile characteristics in adulthood (*53*). The Axolotl remains in its highly regenerative larval stage unless ambient water supply ceases (*54*), the cave-dwelling Olm never leaves its larval stage and is predicted to live >175 years (*55*), and ‘immortal jellyfish’ of the genus *Turritopsis* manage to revert their sexually mature medusa stage into the budding polyp at any time, thus considered to have an indefinite lifespan (*56*). Even amongst these extreme cases humans are considered as neotenic apes due to traits such as orthognathy, body hair reduction, high relative brain weight and prolonged growth period. A remarkable increase of reproductive success with age (*57*) is just one of 43 neotenic traits listed (*52*). Splenic erythropoiesis and expansion of medullary granulopoiesis along with compression of the BC compartment are clear neotenic traits seen in NMRs. It is believed that neoteny is linked to longevity in Axolotl, Olm and human (*52*). Hence, the multiple neotenic features of the hematopoietic system we identified are likely linked to NMR longevity.

**Fig. 8.**
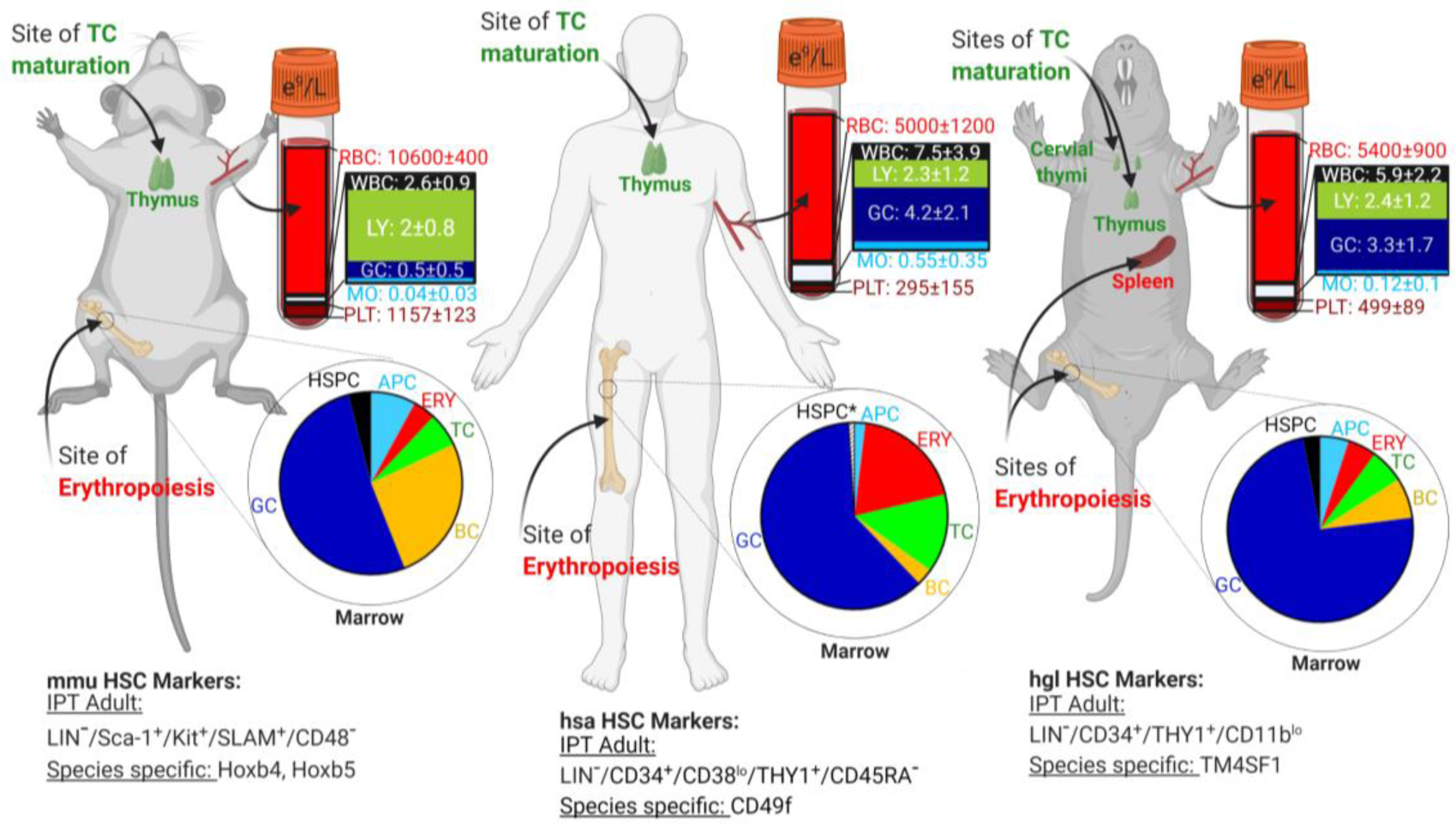
Interspecies differences and similarities in blood and marrow composition. Quick reference including blood and marrow compositions, sites of TC maturation (*11*) and erythropoiesis, and HSC markers. Cell type proportion in bone marrow of mouse and NMR based on Fig 5b.

A striking feature of NMR stem cells was a higher proportion of cells in quiescence. The dynamic equilibrium between quiescence and specific cell cycle kinetics is a hallmark of adult stem cells, hence mouse HSCs have been shown to contain a dormant fraction of ∼20% (*58*). Remarkably we found no significant differences in the frequencies of LTCs, LSKs and human CD34^+^/CD38^lo^ HSPCs in normal BM (Fig. S8c), suggesting conserved stem cell pool frequencies. Our data suggest an expanded quiescent HSC pool in NMRs and rigid control of cell cycle genes at the transcriptional level (Fig. S1h, 2d, 3d). The enlarged quiescent HSC pool would benefit longevity by minimizing damage to stem cells and decelerating clonal expansion, which is a key feature of an aged hematopoietic system (*59*).

Strikingly, CITE-CD11b is correlated with loss of B-lymphopoiesis in xenografts, supporting the conclusion that LTCs transition into CD11b^+^/Thy1.1^int^/CD34^hi^ myeloid- primed CP1. FACS quantitation revealed >4-fold increase of CP1 over LTCs in marrow and >3-fold in spleen (Fig. 2b-f), underscoring a myeloid differentiation bias in NMR hematopoiesis. BM scRNA-Seq revealed 2.5-fold increase of HSPCs in 11 year versus 3 year old NMRs. The common HSPC cluster comprised LTC/CP1/CP3, and we showed that myeloid CP1 progenitors and LTCs increase with age, signs of clonal hematopoiesis in NMRs. However, while an oligopotent myeloid progenitor and the primitive stem cell compartment expand, there is no age-associated increase in PB-WBCs. Moreover, the megakaryocytic differentiation bias, a common hallmark of aging hematopoietic lineage trajectories (*60*), was not as evident in 11 year old NMRs as in 12 month old mice (Fig. 5g-h). Remarkably, blood platelets did not elevate significantly in aged NMRs.

Maintenance of youthful effector cell compositions despite myeloid progenitor expansion in middle-aged animals could be a direct result of an enlarged quiescent HSPC fraction in concert with a prolonged cell cycle, delaying peripheral manifestation of clonal hematopoiesis.

In summary, the entire hematopoietic system of naked mole-rats evolved a combination of unique or neotenic adaptations to an extended healthspan, such as diminished platelets and delay of age-associated leukocytosis, active hematopoiesis in the spleen, and as described in an accompanying report, additional cervical thymi and absence of thymic involution (*11*). On the molecular level all hematopoietic cells feature a slower G1-S-transition, stem and progenitors are less metabolically active than those from short-lived mice and the HSC compartment contains a higher fraction of quiescent cells. NMRs have evolved extreme longevity and resistance to almost all age-related diseases. Understanding the molecular mechanisms of these evolutionary adaptations can lead to novel strategies improving human health. Our resource provides a platform for using NMRs as a research model in stem cell biology, immunology, inflammation and the studies of systemic factors in aging.

## Materials and Methods

### Animals

All animal experiments were approved and performed in accordance with guidelines numbers 2009-054 (naked mole rat) and 2017-033 (mouse). Naked mole rats were from the University of Rochester colonies, housing conditions as described (*61*). C57BL/6 mice were obtained from NIA, in comparative assays yLSK were sorted from 3-4 month and oLSK from 25 month old mice. Immunodeficient strain NSGS [NOD.Cg-*Prkdc^scid^ Il2rg^tm1Wjl^* Tg(CMV-IL3,CSF2,KITLG) 1Eav/MloySzJ] was purchased from JAX.

### Primary cell isolation

Marrow from mice and naked mole-rats was extracted from femora, tibiae, humeri, iliaci and vertebrae by crushing. Spleen, liver, thymus and lymph nodes were minced over a 70µm strainer and resuspended in FACS buffer. Blood from mice was drawn via retroorbital capillary bleeding, naked mole-rat blood was obtained via heart puncture.

Human marrow was obtained from the URMC Pathology and Laboratory Medicine in accordance to RSRB STUDY00006161. Human BM cell fractions shown in Fig. 8 were based on Wintrobe’s monograph (*62*) and cross referenced with Osgood et al (*63*); human marrow HSC fraction was approximated accordingly (*44*).

### Hematology Analyzer

PB parameters were measured with a Vet ABC Plus+ (scil) Analyzer. Specifically, naked mole-rat and mouse samples were measured with the “mouse_research” protocol (scil Tech Support, available upon request), which provides a 3-part differential in 17 parameters.

### Histology

Imaging and analysis was performed using a using a Nikon Eclipse Ti-S microscope. Coverslips were applied with DEPEX Mounting media (Electron Microscopy Sciences), except for Alkaline Phosphatase staining where Vectashield Hard Set Mounting Medium for Fluorescence (Vector) was applied. Femur bones were decalcified with 14% EDTA for a minimum of 2 weeks and stored in 10% neutral buffered formalin. Soft tissues were stored in 10% neutral buffered formalin, processing was done using a Sakura Tissue-Tek VIP 6 automated histoprocessor, paraffin embedding was done using a Sakura Tissue-Tek

TEC 5 paraffin embedding center. A Microm HM315 microtome was used to section tissues at a thickness of 5µm, which then were floated onto a slide with a water bath at a temperature between 45°C and 55°C. Sections were deparaffinized and rehydrated to distilled water through xylene and graded ethanol (100% to 70%).

#### May-Grünwald-Giemsa

Cytospins of whole spleen or WBM or sorted cells were prepared using a Rotofix 32A (Hettich) and stained at room temperature with May-Grünwald solution (Sigma) for 5min, washed in phosphate buffer pH 7.2 (Sigma) for 1.5min, and counterstained in 4.8% Modified Giemsa (Sigma) for 13min.

#### Alkaline Phosphatase

Cytospins were stained with the Alkaline Phosphatase kit (Sigma) according to manufacturer’s instructions with the exemption of combining FBB-Alkaline Solution with Hematoxylin Solution, Gill No. 3 (Sigma) as counterstain.

#### Benzidine Mayer’s Hematoxylin

Slides were fixed at room temperature with methanol for 30sec, incubated with o-Dianisidine (Sigma) 1% in methanol for 1min and stained with H2O2 2.5% in ethanol for 30sec before rinsing for 15sec in water and counterstaining with Mayer’s Hematoxylin Solution (Sigma) for 2min.

#### Wright Giemsa

Blood films were incubated with Wright-Giemsa Stain (Electron Microscopy Sciences) for 1min, rinsed briefly with water and developed in phosphate buffer pH 7.2 for 2min, then rinsed again. Slides were scored by taking 3 random micrographs of monolayers from the feathered edge of each sample to count both RBCs and platelets and average the technical replicates. Then mean RBC levels from the bloodcounter measurements (mouse, 9.1e^12^/l; naked mole-rat 5.4e^12^/l) were used to convert PLT/RBC ratios to a volumetric PLT count via bloodcounter by

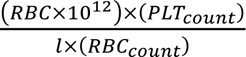

#### Hematoxylin & Eosin

Sections were stained with Mayers Hematoxylin (Sigma) for 1min and washed with tap water to remove excess blue coloring. Soft tissue sections were further decolorized with 3 dips in 0.5% acid alcohol and washed in distilled water. The nuclei of sections were blued in 1X PBS for 1 minute and washed again in distilled water. An Alcoholic-Eosin counterstain was applied for 30sec before slides were immediately dehydrated and cleared through 3 changes of 95% ethanol, 2 changes of 100% ethanol, and three changes of Xylene for 1min each.

#### Microwave Giemsa for plastic marrow sections

Paraffin-embedded Femora were subjected to the microwave modification of a conventional Giemsa stain, which we found to produce clearer contrast of megakaryocytic cells as distinguished by their pale purple cytoplasm and abundant nuclear chromatin staining due to polyploidy. The stain was performed as described in www.urmc.rochester.edu/urmc-labs/pathology. Slides were scored by taking 3 random micrographs of marrow from medullary canal for each sample to count polyploidy giant Megakaryocytes, average the technical replicates and convert micrograph pixel size via magnification to bone area in mm^2^.

### Methylcellulose colony assays

Fresh sorted or whole BM naked mole-rat cells were tested to grow in mouse (M3434, SCT), rat (R3774, SCT) or human (H4435, SCT or HSC005, RnD Systems) methylcellulose formulations to show the highest colony numbers, colony sizes and cell viability with human cytokine cocktails. Either 1x10^4^ whole marrow or 1x10^3^ sorted naked mole-rat cells were added to 3ml of HSC005 supplemented with 1% Penicillin/Streptomycin and 1X GlutaMAX (both Thermo Fisher), equally divided into two 35mm dishes, grown for 21d at 32°C, 5% CO2 and 3% O2, and scored. Although hematopoietic naked mole-rat cells will grow at 37°C, the total number as well as colony and cell type diversity is strongly enhanced at 32°C (data not shown). Colony assays grown at 37°C give rise to two types of colonies (erythroid vs myeloid), which are notably smaller than each of the 4 colony types we can distinguish at 32°C (Fig. S5b). Replatings were done by resuspending scored dishes at day 21 in FACS buffer, count cells and replate 1x10^4^ cells into above growth conditions. CP3 cells did not grow substantially in the first replating, and a second replating had no sizable colonies for all naked mole-rat HSPC types. For Benzidine staining of naked mole-rat methylcellulose assays a 0.2% benzidine dihydrochloride (Sigma) solution in 0.5M acetic acid was prepared, which was supplemented with 0.24% of 50% H2O2. 1ml of this solution was layered carefully over each dish, and past 5min colonies were scored for the proportion of colonies which are uniformly benzidine-unreactive (color-less), uniformly benzidine-reactive (blue) and colonies containing both reactive and unreactive cells (mixed colonies containing both differentiated, hemoglobin containing and non-erythroid cells).

### Seahorse Assay

Sorting for human (Fig. S8a), mouse (Fig. S8b) and naked mole-rat (Fig. 2a-c) marrowSeahorse XF96 Cell Culture Microplates (Agilent Technologies) between 50-250*10^3^ cells per 200µl of the following culture media: Human LT-HSCs (LIN^−^/CD38^lo^/CD34^+^/CD45RA^−^/CD90^+^) in StemSpan Serum-free expansion medium (SFEM; Stemcell Technologies) supplemented with 100ng/ml human SCF, 100ng/ml human FLT3L, 20ng/ml human IL-6, 50ng/ml human TPO (all Peprotech), 0.75µM Stemregenin (SR-1; Stemcell Technologies Cat# 72342), modified as described (*64*). Human MPPs (LIN^−^/CD38^lo^/CD34^+^/CD45RA^−^/CD90^−^) in StemSpan supplemented with 50ng/ml hSCF, 50ng/ml hFLT3L, 10ng/ml hIL-3, 10ng/ml hIL-6, 20ng/ml hTPO (all Peprotech), 0.25% Chemically defined lipid concentrate (CDLC; ThermoFisher Cat# 11905031). Human oligopotent progenitors (hOPP; LIN^−^/CD38^hi^/CD34^+^) in RPMI with 10% FBS (both Gibco), 1% GlutaMAX^TM^ (Thermo Fisher), 5ng/ml hSCF, 5ng/ml hGM-CSF, 5ng/ml hIL-3. Mouse LT-HSCs (LIN^−^/Sca-1^+^/Kit^+^/CD48^−^/SLAM^+^) in the long-term HSC expansion cocktail (*65*). Mouse MPPs (LIN^−^/Sca-1^+^/Kit^+^/CD48^+^/SLAM^−^) in StemSpan with 1% GlutaMAX^TM^, 10ng/ml mSCF, 20ng/ml mTPO (all Peprotech), 10ng/ml mFGF1 (all Peprotech), 20ng/ml mIGF2 (BioLegend Cat# 588204). Mouse LIN^−^/Kit^+^/Sca-1^−^ (LK; mOPP) in StemSpan with 10% FBS, 10ng/ml mSCF, 10ng/ml mIL-3, 10ng/ml mIL-6 (all Peprotech). Naked mole-rat LTC (LIN^−^/Thy1.1^int^/CD34^hi^; CP2) in StemSpan with 1% GlutaMAX^TM^, 1% CDLC, 100ng/ml hSCF, 100ng/ml hFLT3L, 20ng/ml hIL-6, 50ng/ml hTPO, 1µM SR-1, 0.1µM UM-171 (Selleck Chemicals Cat# S7608). Naked mole-rat CP1 (LIN^+^/Thy1.1^int^/CD34^hi^) in StemSpan with 1% GlutaMAX^TM^, 1% CDLC, 50ng/ml hSCF, 50ng/ml hFLT3L, 20ng/ml hIL-6, 10ng/ml hGM-CSF, 1µM UM-729 (Stemcell Technologies Cat# 72332). Naked mole-rat CP3 (LIN^−^/Thy1.1^lo^/CD34^hi^; MEP) in StemSpan with 1% GlutaMAX^TM^, 1% CDLC, 50ng/ml hSCF, 50ng/ml hTPO, 1 U/ml hEPO (all Peprotech). All expansion cocktails were added with 1% Penicillin-Streptomycin (Thermo Fisher Cat# 15140163). Cells were allowed to settle for 16-20h at 37°C (32°C for Naked mole-rat), 5% CO2 and 0.5% O2. We used Corning Cell-Tak Cell and Tissue Adhesive (Thermo Fisher Cat# CB-40240) at 22.4µg/ml concentration per well to prepare Cell-Tak coated XF96 microplates according to the manufacturers guidelines (Agilent Technologies). Cells were seeded into the coated microplates immediately before the assay by centrifugation with 200g for 1min without brake. Subsequently we strictly adhered to the Seahorse XF Cell Mito Stress Test Kit protocol (Agilent Technologies Cat# 103015-100). Cells were counted before and after the assay using a Celigo S Image cytometer (Nexcelom Biosciences) with automated 96-well Brightfield imaging at the

Technologies Cat# 103681-100). Final Well concentrations were 1.5µM Oligomycin, 1µM FCCP and 0.5µM Rotenone/Antimycin A. Measurements were taken on a Seahorse XFe96 Analyzer in the URMC Flow core using Wave 2.6.1 software (Agilent Technologies).

### Flow Cytometry

<u>Flow cytometry analysis</u> was performed at the URMC Flow Core on a LSR II or LSRFortessa (both BD), or on our labs CytoFlex S (Beckman Coulter). Kaluza 2.1 (Beckman Coulter) was used for data analysis. Staining and measurement were done using standard protocols. Red blood cell lysis was done by resuspending marrow pellets in 4ml, spleen pellets in 1ml and up to 500µl blood in 20ml of RBC lysis buffer, prepared by dissolving 4.1g NH4Cl and 0.5g KHCO3^−^ into 500ml double-distilled H2O and adding 200µl 0.5M EDTA. Marrow and spleen were incubated for 2min on ice, blood was lysed for 30min at room temperature. Cells were resuspended in FACS buffer (DPBS, 2mM EDTA, 2% FBS [Gibco]) at 1x10^7^ cells/ml, antibodies were added at 1µl/10^7^ cells, vortex-mixed and incubated for 30min at 4°C in the dark. DAPI (Thermo Fisher) @ 1µg/ml was used as viability stain. The primary gating path for all unfixed samples was: scatter-gated WBC (FSC-A vs SSC-A) => singlets1 (SSC-W vs SSC-H) => singlets2 (FSC-W vs FSC-H) => viable cells (SSC vs DAPI) == proceed with specific markers/probes. Compensation was performed using fluorescence minus one (FMO) controls for each described panel. For antibody validation we incubated 1mio cells in 100µl Cell Staining Buffer (BioLegend; Cat# 420201) and added 5µl Human TrueStain FcX™ and 0.5µl TruStain FcX™ PLUS, followed by incubation for 10min at 4°C. We then proceeded withfluorescent antibody staining as above.

<u>Immunophenotyping</u> of naked mole-rat BM, spleen, thymus, PB and lymph nodes: CD90 FITC; CD125 PE; Thy1.1 PE-Cy7; CD34 APC, CD11b APC-Cy7. Quantification of murine BM SLAM HSCs was performed using mouse LIN Pacific Blue; Sca-1 BUV395; CD150 PE; Kit PE-Cy7; CD48 APC-Cy7. Quantification of human BM LT-HSCs was performed using human LIN Pacific Blue; CD34 APC; CD38 APC-Cy7; CD45RA FITC; CD90 PE-Cy7. Fluorescence minus one (FMO) controls were applied for fluorescent spillover compensations for each species and tissue used. All antibodies can be found in Table S7.

<u>Sorting</u> was performed at the URMC Flow Core on a FACSAria (BD) using a 85μm nozzle, staining was done as described. Human HSCs were sorted for population RNA-Seq as LIN^−^/CD34^+^/CD38^Lo^/CD45RA^−^/CD90^Dim^ (Fig. S5A). Naked mole-rat HSPC populations were sorted as described with a lineage cocktail comprised of CD11b, CD18, CD90 and CD125 (NMR LIN). Naked mole-rat marrow and spleen sorting panel was: NMR LIN Pacific Blue; Thy1.1 PE-Cy7; CD34 APC. Naked mole-rat blood sorting panel was: Thy1.1 PE-Cy7; CD11b APC-Cy7.

<u>Molecular probing</u> was performed on frozen aliquots from mouse and naked mole-rat BM. For each probe, cells were diluted in 1ml pre-warmed DMEM+ at 1x10^6^ cells/ml. All stainings were performed simultaneously for 4-6 naked mole-rat, 2-4 old, 2-4 young mice and 4 human biological replicates. <u>ALDEFLUOR</u> (SCT) reagent was added at 0.5µl/ml, mixed and incubated for 15min at 37°C in a water bath. <u>MitoStatus TMRE</u> (BD) was added to 0.5x10^6^ cells/ml with 25nM and incubated for 10min at room temperature in the dark. FCCP (Trifluoromethoxy carbonylcyanide phenylhydrazone) was added to negative controls at 5µM during TMRE staining. <u>JC-1</u> (Thermo Fisher) was added at 1µM with or without 5µM FCCP and incubated for 15min at 37°C. <u>MitoTracker Orange CMTMRos</u> (Thermo Fisher) was added at 10nM and incubated for 45min at 37°C. <u>MitoSOX red</u> (Thermo Fisher) was added at 5µM and incubated for 30min at 37°C. CellROX <u>Orange</u> (Thermo Fisher) was added at 5µM and incubated for 60min at 37°C. The subsequent antibody staining was performed as above with 30min incubation on ice, panel was Sca-1 (mouse) or CD34 (naked mole-rat) APC; Kit (mouse) or Thy1.1 (naked mole-rat) PE-Cy7; Lineage Cocktail V450. <u>Rhodamine 123</u> staining was performed by incubating 1x10^6^ cells for 30min with 1µg/ml Rho in HBSS+ (HBSS, 2% FBS, 10mM HEPES; all Gibco) at 37°C, then cells were washed with 2ml HBSS+, spun down and reincubated for 15min at 37°C.

### Pyronin Y staining

Mouse and naked mole-rat BM cells from frozen aliquots were count-adjusted to 1x10^6^ cells/ml and resuspended into 1ml of DMEM+ (DMEM high Glucose, 2% FBS, 10mM HEPES; all Gibco). Upon addition of 50μg/ml Verapamil (Sigma) and 5μM DyeCycle Violet (Thermo Fisher) cells were incubated for 45min at 37°C in a water bath, vortex-mixed every 15min. Past 45min 0.1μg/ml Pyronin Y was added to the reaction and (HBSS [Gibco], 0.33M HEPES, 3.5% FBS, 0.02% NaN3 [Sigma]). A subsequent antibody staining was performed as above with incubation on ice, panel was Sca-1 (mouse) or CD34 (naked mole-rat) APC; Kit (mouse) or Thy1.1 (naked mole-rat) APC-Cy7; Lineage Cocktail FITC; 500nM SYTOX Green (Thermo Fisher) was used as viability stain.

### Ki67 staining

Mouse and naked mole-rat BM cells from frozen aliquots were count-adjusted to 1x10^7^ cells/ml and antibody staining was performed as described, panel was Sca-1 (mouse) or CD34 (naked mole-rat) APC; Kit (mouse) or Thy1.1 (naked mole-rat) APC-Cy7; Lineage Cocktail FITC. For fixation and permeabilization we used the buffers from the BrdU Flow Kit (BD). Briefly, cells were fixed for 30min in Cytofix/Cytoperm on ice at 100µl/1x10^6^ cells, permeabilized for 10min in CytopermPlus on ice at 100µl/1x10^6^ cells, refixed for 5min in Cytofix/Cytoperm on ice at 100µl/1x10^6^ cells, all washes done with 1X Perm/Wash. Cells were resuspended in Staining buffer at 1x10^7^ cells/ml, Ki67 antibodies (mouse: clone 16A8; naked mole-rat: clone Ki-67; both PE-conjugated, BioLegend) were added at 5µl/1x10^6^ cells and incubated for 30min at room temperature in the dark, 1µg/ml DAPI was used as DNA stain.

### EdU-BrdU dual-Pulse labelling

Mice aged 6 months or naked mole-rats aged 2-4 years were intraperitoneally (i.p.) injected with 1mg (2′S)-2′-Deoxy-2′-fluoro-5-ethynyluridine (F-ara-EdU; Sigma) from a 10mg/ml stock in DMSO diluted with sterile 0.9% sodium chloride solution (Sigma).

Exactly 2h later animals were i.p. injected with 2mg 5-Bromo-2′-deoxyuridine (BrdU; Sigma) from a 20mg/ml stock in DMSO diluted with sterile 0.9% sodium chloride solution (Sigma). Animals were euthanized for tissue harvest 30min post BrdU administration. Mouse and naked mole-rat BM cells from frozen aliquots were count-adjusted to 1x10^7^ cells/ml. Antibody staining was performed as described before fixation, panel was LIN-V450/BV421, Sca-1 (mouse) or CD34 (naked mole-rat) APC, Kit (mouse) or Thy1.1 (naked mole-rat) APC-Cy7. We used the fixing and permeabilization buffers from the Click-iT EdU Plus Kit (Thermo Fisher). Antibody-stained cells were washed twice in PBS 1% BSA (Cell Signaling Technology), then resuspended with 100µl/1x10^6^ cells Fixative and incubated for 15min at room temperature (RT) in the dark. cells were 100µl/1x106cells Perm/Wash buffer and incubated for 15min at RT in the dark. Click-iT Plus reaction cocktail was prepared according to the Kit (Thermo Fischer), directly added to the permeabilization mix and incubated for 30min at RT in the dark. Cells were washed two times with Perm/Wash and resuspended in 100µl of 300µg/ml DNAse1 into 30µg/1mio cells, and incubated for 1h at 37°C in a waterbath. Cells were washed with Perm/Wash and stained with 1µl/1mio cells anti-BrdU from the FITC BrdU Flow Kit (BD Biosciences) for 20min at RT in the dark. For EdU cell cycle measurements no DNAse1 digestion and BrdU labelling was performed, instead cells were stained with 500nM SYTOX Green.

### Xenotransplantations

Naked mole-rat BM and/or spleen cells were extracted, sorted and directly transplanted into 2.5Gy-irradiated (24h pre Tx) NSGS recipients between 5-9 weeks of age at cell doses between 50-100k sorted or 1-5mio whole marrow naked mole-rat cells. Injections were done via the retroorbital sinus, blood sampling was performed via maxillary vein or retroorbital plexus at weeks 4, 8 and 12. Hosts were culled at 2, 4, 8 or 12 weeks and engraftment frequencies were estimated by flow cytometry using only naked mole-rat markers not cross-reactive with mouse cells and CD45.1 (A20, BioLegend). Engraftment rates were adjusted for input cell dose to 100k/Tx. Gating path was WBC (FSC-A vs SSC-A) => singlets1 (SSC-W vs SSC-H) => singlets2 (FSC-W vs FSC-H) => viable cells (SSC vs DAPI) => NOT Thy1.1^−^/CD34^−^ (CD34 vs Thy1.1) == engrafted naked mole-rat cells (Fig. 4d). One limitation for quantifying engraftment levels is that naked mole-rat BM features cells negative for the above markers which can arise from transplanted HSPCs as xenogenic CP7 (Fig. 4d). A cross-reactive guinea pig CD45 antibody does not stain >80% of naked mole-rat WBM cells and exhibits notable cross-reactivity with BM from NSGS recipients (Fig. S6g-j). We further detected cells double-positive for guinea pig CD45 and CD45.1. Cells stained as Thy1.1^+^ and/or CD34^+^ are clearly originated by the xenograft as untransplanted NSGS BM does not feature any Thy1.1 of CD34 labelled cells (Fig. 4d). Since all three different cell populations from guinea pig CD45 vs CD45.1 staining (DN, CD45.1^+^, CD45^+^/CD45.1^−^) contain a different pattern of cells stained as Thy1.1^+^ and/or CD34^+^, we considered any cell positive for one or both markers as xenograft. We reasoned that due to the *in vitro* cross-reactivity of human SCF engraftment would be supported when using NSGS hosts. However, when we compared the engraftment efficiency for ∼1x10^5^ LTCs transplanted into NSGB (NOD.Cg-*B2m^tm1Unc^ Prkdc^scid^ Il2rg^tm1Wjl^*/SzJ) or NSGS mice at 4 weeks and same cell dose between NSG (NOD.Cg-*Prkdc^scid^ Il2rg^tm1Wjl^*/SzJ) and NSGS at 8 weeks, we found no significant differences between the strains (data not shown).

### 5-FU treatments

Mice aged 6 months or naked mole-rats aged 2-4 years were given intraperitoneal (i.p.) injections with 150mg/kg 5-Fluorouracil (5-FU; Sigma) from a 50mg/ml stock in DMSO diluted with sterile 0.9% sodium chloride solution (Sigma). Animals were monitored daily and euthanized moribundity.

### Quantitative PCR

Mouse and Naked mole-rat sorted TCs and thymic tissue were used for RNA extraction by Trizol (Thermo Fisher). RNA was quantified using a NanoDrop One (Thermo Fisher), and 100ng was used as input for the High Capacity cDNA Reverse Transcription Kit (Thermo Fisher). RT reaction was performed according to instructions and the 20µl reaction diluted to 200µl, of which 5µl were used per qPCR reaction. We used iTaq Universal SYBR Green Supermix (Bio-Rad) on a CFX Connect® RealTime System (Bio-Rad) with a three-step cycling of 10sec 95°C, 20sec 60°C, 30sec 72°C for 40 cycles. All primers (IDTDNA) were validated to amplify a single amplicon at the above PCR conditions by gel electrophoresis. Gene sequences for primer design by Primer3Plus were retrieved from ENSEMBL, with the exception of the T cell receptor C-region genes for naked mole-rat. Here we used the WBM RNA-Seq from the transcriptome assembly below to map those genes in a recently published naked mole-rat genome (*66*) using Apollo software and custom scripts. For absolute copy number quantitation, qPCR amplicons were gel-purified using the QIAquick Gel Extraction Kit (Qiagen) and subcloned into the pCR2.1 plasmid using the TOPO-TA cloning Kit (Thermo Fisher). Plasmids were prepared using the QIAprep Spin Miniprep Kit (Qiagen). Sanger sequencing was performed by Genewiz using M13 forward and reverse primers. Standard curves were prepared across a 10-fold dilution range from 20ag to 20pg of plasmid DNA. All amplicon gel images, amplicon plasmids and standard curve data is available upon request, Primers can be found in Table S7.

### Transcriptome assembly

All Naked mole-rat RNA-Seq was performed with the [GRC URMC Rochester](https://www.urmc.rochester.edu/research/rochester-genomics-center.aspx). RNA from whole bone marrow (WBM) was sequenced with ∼230 million reads on a HiSeq2500v4 (Illumina). Raw Illumina paired-end sequencing reads where assessed with FastQC. Rcorrector was used to correct sequencing errors and read pairs with uncorrectable errors were removed using a custom python script (GRC URMC Rochester). Adapter and base quality trimming was performed using trim galore and cutadapt resulting in high quality reads that were used as input to Trinity for assembly. FRAMA (*14*) was used to post-process the *de novo* assembly, including reduction of contig redundancy, ortholog assignment using human as a reference, correction of misassembled transcripts, scaffolding of fragmented transcripts and coding sequence identification. Quality assessment of the final FRAMA transcriptome was performed using BUSCO and TransRate. The transcriptome was mapped by blastn to the naked mole-rat genome (hetgla_female_1.0) or to transcript sequences annotated in ENSEMBL97. Mapped genomic coordinates of transcripts were thus compared to those of annotated genes using a custom python script. We found that 512 non-overlapping FRAMA transcripts (i.e., gene loci) were absent from the annotation, and another 5281 had >20% transcript length mapped to the genome but not matching annotated isoforms (Table S1).

### Population RNA-Seq

RNA from sorted human and naked mole-rat populations was sequenced at ∼100 million reads on a HiSeq2500v4 (Illumina), the SMARTer® Ultra® Low RNA Kit (Takara) was used for library preparation. All GEO datasets for human and mouse HSPC populations were acquired with SRA toolkit and processed from raw fastq files. Raw Illumina paired-end sequencing reads where subjected to base quality trimming using Trimmomatic and were assessed with FastQC. *RSEM v1.3.0* with *STAR* aligner option was used to calculate expected counts and TPMs (*67*). We used a customized perl script to run RSEM with the FRAMA transcriptome as reference using *bowtie2*aligner option. We also run RSEM with the ENSEMBL94 hetgla_female_1.0 annotation using *STAR* aligner option to confirm all clusterings and differential gene expression signatures for all naked mole-rat samples, results were almost identical to those obtained with FRAMA (data not shown).

Subsequent analysis was done with *R 4.0.2* and *Bioconductor* (*68*). Expected counts from different transcript isoforms of the same gene were added up to one unique identifier (uniquefy) using ddply and numcolwise functions of the *plyr* package, *edgeR* was used to calculate size factors with method=”RLE” and computing CPMs. We applied *genefilter*to calculate the interquartile range (IQR) of CPMs with IQR(x) > 1 to filter unexpressed and outlier genes; library-size normalized, IQR-filtered log2-transformed CPMs were vst-transformed by *DESeq2*, then a PCA from stats package was used as input for *Rtsne*. We applied *limma* to perform voom-transformation and select for differentially expressed genes (DEGs) with p < 0.05 and log-fold-change 1.

GSEA was performed using the *gsva* package with method=”ssGSEA” using either the hematopoietic stem and progenitor geneset collection modified from Schwarzer & Emmrich et al (*69*) in Table S1 or the MSigDB v6.0 hallmark genesets with a p-value threshold of 0.05. All GSEA calculations were performed on the combined up- and downregulated DEG signature for each group, see Table S4. The *fGSEA* package was used to retrieve leading edge genes after reperforming GSEA under default conditions (*70*), the required gene rank metric was generated according to (*71*). All naked mole-rat population RNA-Seq DEG signatures (Table S4) were used to create genesets and were added to Table S1.

The expression gradient in Fig. 4b was calculated by a customized R function, which ordered the log2-transformed CPMs for each gene along their numeric value, allowing to filter out the genes subsequently changing expression from one group to another, see Table S4.

For the 3-species comparison uniquefied human, mouse and naked mole-rat TPM datasets (Fig. 6a, Fig. S8d-e) were merged based on HGNC symbols, then *genefilter* was used to calculate IQR of TPMs with IQR(x) > 1 to filter unexpressed and outlier genes; The *TCC* package was used to calculate TMM-based size-factors. The function betweenLaneNormalization with median scaling from the *EDASeq* package was used to normalize for sequencing batch effects. We used the *RUVSeq* package to normalize TPMs for batch effects across datasets. The *limma* package was used to plotMDS of the full 3- species dataset (Fig. S8d), see Table S6 (sheet “metadata.population.RNA-Seq”). Next we split the dataset collection into three subsets based on developmental stage of each population. TPMs were vst-transformed by *DESeq2*, then a PCA from the *stats* package was used as input for *Rtsne* (Fig. S8e); DGE and GSEA were performed as above using the population species as contrast and the MSigDB v6.0 hallmark genesets.

### Single cell RNA-Seq

#### Naked mole-rat sorted CITE-Seq data (Fig. 1, Fig. S1)

Marrow, blood and thymus cells from 2 animals aged 11m (♀&♂) were enriched by sorting. For BM we sorted CP1 3k, LTC 3k, CP3 2k, CP4 2k, CP5 2k, CP6 2k, CP7 3k, LIN^+^/CD34^−^ 1.5k, LIN^+^/CD34^+^ 1.5k; total 40,000 marrow cells from 2 animals as three 10X v2 chemistry libraries (2 replicates LIN^−^ pooled, one replicate LIN^+^ pooled; Fig. S1E). For PB we sorted GC 1.5k, MO 1k, BC 1k, TC 1.5k; total 10,000 peripheral blood leukocytes from same animals as above into one pooled 10X v2 chemistry library (Fig. S1D). For thymus we sorted CP8 1k, CP9 1k, LTC 1k; total 6,000 thymocytes from same animals as above into one 10X v2 chemistry library (Fig. S6H). Cells were pooled according to their tissue origins and processed for CITE-Seq using a [protocol](https://citeseq.files.wordpress.com/2019/02/cite-seq_190213.pdf) from the Stoeckius lab and the Chromium Single-Cell 3′ Library & Gel Bead Kit v2 (10X Genomics)(*13*). Raw reads generated on the Illumina NovaSeq6000 sequencer were demultiplexed using *[Cellranger](https://support.10xgenomics.com/single-cell-gene-expression/software/pipelines/latest/what-is-cell-ranger) 3.0.2* software in conjunction with Illumina’s *bcl2fastq 2.19.0*. *Cellranger* was also used to align the read data to the FRAMA *de novo* transcriptome assembly and ENSEMBL94 hetgla_female_1.0, barcode count, UMI compress, and filter for “true” cells. CITE-Seq data for each capture was also demultiplexed using *bcl2fastq*and processed with *CITE-seq-Count 1.4.2* (*72*) given the antibody barcode sequences, a white list of filtered cell barcodes from the matching Cell Ranger “count” run, and parameters: “-cbf 1 -cbl 16 - umif 17 -umil 26”.

#### Subsequent analysis was done with *R 4.0.2* and *Bioconductor*

The marrow and blood libraries were merged and FRAMA Trinity isoforms were uniquefied by row-wise addition of UMI-counts for each isoform of the same gene using a *data.table* snippet.10X files were assigned to a *SingleCellExperiment* S4 class, and each gene without any counts in any cell was removed. We converted the S4 class into a *Seurat 3.1* object and added the CITE-signals in form of an independent “assay”, barcodes were quality filtered to keep cells between 200-5,000 detected genes/cell and <25,000 counts per cell. RNA assay was log-normalized with “scale.factor = 1e4”, CITE assay was “CLR” normalized. Variable features were detected with arguments selection.method = “vst”, nfeatures = 3000. Scores respective *Seurat* [vignette](https://satijalab.org/seurat/v3.1/cell_cycle_vignette.html). Clustering was done using *Seurat’s* FindClusters function with resolution = 0.5. Next we used the doublet detection and removal workflow as suggested in the *[Bioconductor OSCA vignette](http://bioconductor.org/books/release/OSCA/)*. Briefly, we run findDoubletCluster from the *scDblFinder* package, followed by *in silico* simulation of doublets from the single-cell expression profiles (*73*) using computeDoubletDensity from *BiocSingular* package, and excluded any cluster which was identified in both methods. The DEGs for each cluster were detected by FindAllMarkers function with arguments test.use = “MAST”, logfc.threshold = log(2), min.pct = 0.25, return.thresh = 0.05. Hematopoietic cell type annotation was done through *fGSEA* using the modified HSPC geneset collection (*69*), extended with the upregulated DEGs from the naked mole-rat population RNA-Seq analysis, upregulated DEGs from joint analysis of murine HSPCs from multiple studies (*74–80*), upregulated DEGs from human HSPC population RNA-Seq datasets (*81–84*), selected genesets from MSigDB and Immgen databases and example genesets from the *[SingCellaR software](https://supatt-lab.github.io/SingCellaR.Doc/)* (Table S1). To determine the rank metrics for *fGSEA* the q-value requires to be transformed by -log10(q-value) (*71*). *Seurat’s* FindAllMarkers function can generate 0 q-values (p_val_adj, Table S2) for high confidence hits, thus for any 0 we added the lowest q-value > 0 of the entire marker list for the group to test to each marker with q-value = 0. This generates ties in the pval ranking by fGSEA for the genes with modified 0 q-values, which are automatically resolved by retaining their order according to their fold-change of expression. A custom script was generated to pipe fGSEA with our HSPC geneset collection (Table S1) through each clusters marker genes, results are deposited in Table S2. The entire process was done in an iterative manner to condense multiple clusters of the same overabundant cell type (e.g. neutrophil granulocytes) into one partition, while maintaining distinctive low abundance clusters. Single cell expression maps (Fig. S1i, 2f, 3f) were done with *schex* package using nbins = dim(Seurat.object)[2]/200. *PhateR* was used as suggested by running an initial graph imputation, and obtaining the final graph with parameters knn=8, decay=100, t=25 (*19*).

#### Human Cell Atlas data (Fig. S2)

The original data comprising 380,000 marrow cells from 8 human donors is available from the [HCA data portal](https://data.humancellatlas.org/explore/projects/cc95ff89-2e68-4a08-a234-480eca21ce79) or as the *HCAData R* package. We used a subset of this dataset available through the *SeuratData* package, randomly downsampled to 40,000 cells. The cell type annotation was obtained by [reference mapping](https://satijalab.org/seurat/articles/multimodal_reference_mapping.html#example-2-mapping-human-bone-marrow-cells-1) according to the *Seurat* vignette. The respective reference was created by weighted nearest neighbor analysis of CITE-Seq data from human marrow according to the *Seurat* [vignette](https://satijalab.org/seurat/articles/weighted_nearest_neighbor_analysis.html) (*40*). Variable features were detected with arguments selection.method = “vst”, nfeatures = 3000. Cell cycle scoring and doublet detection were performed as described above.

Clustering and marker gene identification was done using the same parameters as for naked mole-rat sorted CITE-Seq data. Hematopoietic cell type annotation was done as described above (Table S2). *PhateR* was run with parameters knn=3, decay=100, t=12.

#### RNAMagnet data (Fig. S3)

We used the processed main dataset together with the prior cell type annotation (*85*). Barcodes of tissue type “bone” were excluded, leaving Kit^+^ HSPCs, WBM and CD45^−^ cells in the dataset. Gene features were uniquefied with *data.table*, variable features were detected with arguments selection.method = “vst”, nfeatures = 2000. Cell cycle scoring and doublet detection were performed as described above. Clustering and marker gene identification was done using the same parameters as for naked mole-rat sorted CITE-Seq data. Hematopoietic cell type annotation was done as described above (Table S2). *PhateR* was run with parameters knn=3, decay=100, t=28.

#### Calico data (Fig. S5)

We downloaded the raw fastq files for mouse and naked mole-rat scRNA-Seq from spleen (*26*) using SRA toolkit. *Cellranger 3.1.0* (10X Genomics) was used to generate reference and count matrices for mouse data from ENSEMBL99 or from FRAMA for naked mole-rat. Barcodes were quality filtered to keep cells between 200- 2,500 detected genes/cell and <10,000 counts per cell. Gene features were uniquefied with *data.table*, variable features were detected with arguments selection.method = “vst”, nfeatures = 2000. Cell cycle scoring and doublet detection were performed as described above. Clustering and marker gene identification was done using the same parameters as for naked mole-rat sorted CITE-Seq data. Hematopoietic cell type annotation was done as described above (Table S3).

#### Xenograft data (Fig. 4m-n, Fig. S6m-n)

Host WBM from frozen stocks were subjected to CITE-Seq using the Chromium Single-Cell 3′ Library & Gel Bead Kit v3 (10X Genomics). Cells were processed for TotalSeq™ CITE reagents according to the [manufacturers instructions](https://www.biolegend.com/en-us/protocols/totalseq-a-antibodies-and-cell-hashing-with-10x-single-cell-3-reagent-kit-v3-3-1-protocol) (BioLegend), using both human and mouse Fc blocking reagents (BioLegend). Following fluorescent antibody staining samples were sorted for xenograft cells by Thy1.1/CD34 staining and excluding the “mouse” gate as shown in Figure 3G. For library preparations see below for 10X v3 chemistry. *Cellranger 3.1.0* was used to generate count matrices from both an ENSEMBL94 mouse reference and FRAMA for the same library. Gene features were uniquefied with *data.table*, barcodes were quality RNA assay was log-normalized with “scale.factor = 1e4”, CITE assay was “CLR” normalized. Variable features were detected with arguments selection.method = “vst”, nfeatures = 3000. Libraries were integrated using FindIntegrationAnchors with dims = 1:50, anchor.features = 3000, reduction = “cca”. Mouse cells were removed by subsetting the integrated *Seurat* object. Cell cycle scoring and doublet detection were performed as described above. Clustering and marker gene identification was done using the same parameters as for naked mole-rat sorted CITE-Seq data. CITE feature/antibody marker detection was done as described for transcript cluster markers with the exception of test.use = “wilcox”. Hematopoietic cell type annotation was done as described above (Table S5).

#### Unfractionated BM data (Fig. 5, Fig. S7)

10,000 DAPI^−^ BM cells from 2 mice aged 3m (♀&♂) and 2 mice aged 12m (♀&♂), or 2 naked mole-rats aged 3yr (♀&♂) and 3 naked mole-rats aged 11yr (♀&♂), were subjected to CITE-Seq using Chromium Single-Cell 3′ Library & Gel Bead Kit v3 (10X Genomics). Cells were processed for TotalSeq™ CITE reagents according to the [manufacturers instructions](https://www.biolegend.com/en-us/protocols/totalseq-a-antibodies-and-cell-hashing-with-10x-single-cell-3-reagent-kit-v3-3-1-protocol) (BioLegend), using both human and mouse Fc blocking reagents (BioLegend). Cellular suspensions were loaded on a Chromium Single-Cell Instrument (10x Genomics, Pleasanton, CA, USA) to generate single-cell Gel Bead-in-Emulsions (GEMs). Single-cell RNA-Seq libraries were prepared using Chromium Next GEM Single Cell 3′ GEM, Library & Gel Bead Kit v3.1 (10x Genomics). The beads were dissolved and cells were lysed per manufacturer’s recommendations. GEM reverse transcription (GEM-RT) was performed to produce a barcoded, full-length cDNA from poly-adenylated mRNA. After incubation, GEMs were broken and the pooled post-GEM-RT reaction mixtures were recovered and cDNA was purified with silane magnetic beads (DynaBeads MyOne Silane Beads, PN37002D, ThermoFisher Scientific). The entire purified post GEM-RT product was amplified by PCR. This amplification reaction generated sufficient material to construct a 3’ cDNA library. Enzymatic fragmentation and size selection was used to optimize the cDNA amplicon size and indexed sequencing libraries were constructed by End Repair, A-tailing, Adaptor Ligation, and PCR. Final libraries contain the P5 and P7 priming sites used in Illumina bridge amplification. In parallel, CITE-seq library amplification is performed following SPRI bead purification of CITE-seq cDNA using Q5 Hot Start HiFi Master Mix (New England Biolabs, Ipswich, MA), SI PCR primer (IDT, Coralville, IA), and indexed TruSeq Small RNA PCR primers (Illumina, San Diego, CA) as specified(*13*). Amplified CITE-seq libraries are purified using AMPure XP (Beckman Coulter, Indianapolis, IN) beads and quantified by Qubit dsDNA assay (ThermoFisher, Waltham, MA) and Bioanalyzer HSDNA (Agilent, Santa Clara, CA) analysis. CITE-seq libraries were pooled with 10x Genomics gene expression libraries for sequencing on Illumina’s NovaSeq 6000. Barcodes were quality filtered to keep cells between 200-5,000 detected genes/cell and <20,000 counts per cell. RNA assay was log-normalized with “scale.factor = 1e4”, CITE assay was “CLR” normalized. Variable features were detected with arguments selection.method = “vst”, nfeatures = 3000. Canonical correlation analysis (CCA) was used to integrate libraries (*40*) from either species with FindIntegrationAnchors with dims = 1:50, anchor.features = 3000, reduction = “cca”. Cell cycle scoring and doublet detection were performed as described above. Clustering and marker gene identification was done using the same parameters as for naked mole-rat sorted CITE-Seq data. CITE feature/antibody marker detection was done as described for transcript cluster markers with the exception of test.use = “wilcox”. Hematopoietic cell type annotation was done as described above (Table S5). Differentially expressed markers between age groups for either species used FindMarkers with test.use = “MAST”, logfc.threshold = log(2), min.pct = 0.1. Next we run *fGSEA* with the MSigDb hallmark geneset as mentioned in population RNA-Seq, and plot any pathway with FDR < 0.05 (Fig. S4G; no significant pathways for naked mole-rat markers across age). Differential abundance (DA), testing the cell abundances for clusters across conditions, was performed as described (*86*). Briefly, *edgeR* was used to apply negative binomial generalized linear model dispersion to each library as outlined in the [OSCA Bioconductor collection](http://bioconductor.org/books/release/OSCA/multi-sample-comparisons.html#differential-abundance). SCTransform was used to integrate scaled, clustered and annotated mouse and naked mole-rat unfractionated BM datasets (*43*): SelectIntegrationFeatures with nfeatures = 3000, FindIntegrationAnchors with normalization.method = “SCT”. Cell cycle scoring, clustering and marker detection performed as described above. Conserved markers were identified by running FindConservedMarkers for each cluster across species of the SCT-integrated dataset with test.use = “MAST”, logfc.threshold = log(2), min.pct = 0.25. Differentially expressed markers per cluster between species run FindMarkers with test.use = “MAST”, logfc.threshold = log(2), min.pct = 0.1. We performed *fGSEA* with the MSigDb hallmark geneset and plot any pathway with FDR < 0.05 (Fig. S4I).

### Quantification and Statistical Analysis

Data are presented as the mean ± SD. Statistical tests performed can be found in the figure legends. P values of less than 0.05 were considered statistically significant. Statistical analyses were carried out using Prism 9 software (GraphPad) unless otherwise stated.

## Acknowledgments

The authors thank Alex Aiezza II for RSEM builds of mouse samples, Anthony Corbett for FRAMA, Jason R Myers for a Perl FRAMA-into-RSEM interface, Cameron Baker for cellranger preprocessing of scRNA-Seq data, Michelle Zanche, Jeffrey Malik and John Ashton for Genomic Research Core support.

## Author contributions

S.E. designed and supervised research, performed most experiments and analyzed data; A.T. wrote scripts for single-cell fGSEA and contributed to bioinformatics analyses and FACS; F.T.Z. performed histology quantifications, animal perfusions and data analysis; X.Z. improved genome assembly; M.M. performed FRAMA-genome alignments; M.E.S. performed histology and most histochemistry stainings; Q.Z. mapped TCR genes; Z.Z. performed mouse blood analysis; M.D.G. performed histology and provided human BM specimen; S.G., E.M.I., Z.K., M.T., J.H.K, Z.Z. and V.N.G. contributed to data analysis; A.S. and V.G. supervised research; S.E., A.S. and V.G. wrote the manuscript with input from all authors.

## Data and materials availability

The RNA-Sequencing data that support the findings of this study are available in figshare.com with the identifiers 10.6084/m9.figshare.c.5472735, 10.6084/m9.figshare.c.5470587, 10.6084/m9.figshare.c.5472684, 10.6084/m9.figshare.c.5474256. The custom code to run GSEA for scRNA-Seq clusters for cell type annotation was deposited in GitHub under https://github.com/alex-trapp/sc-fgsea.

## Supplementary Materials

**Fig. S1.**
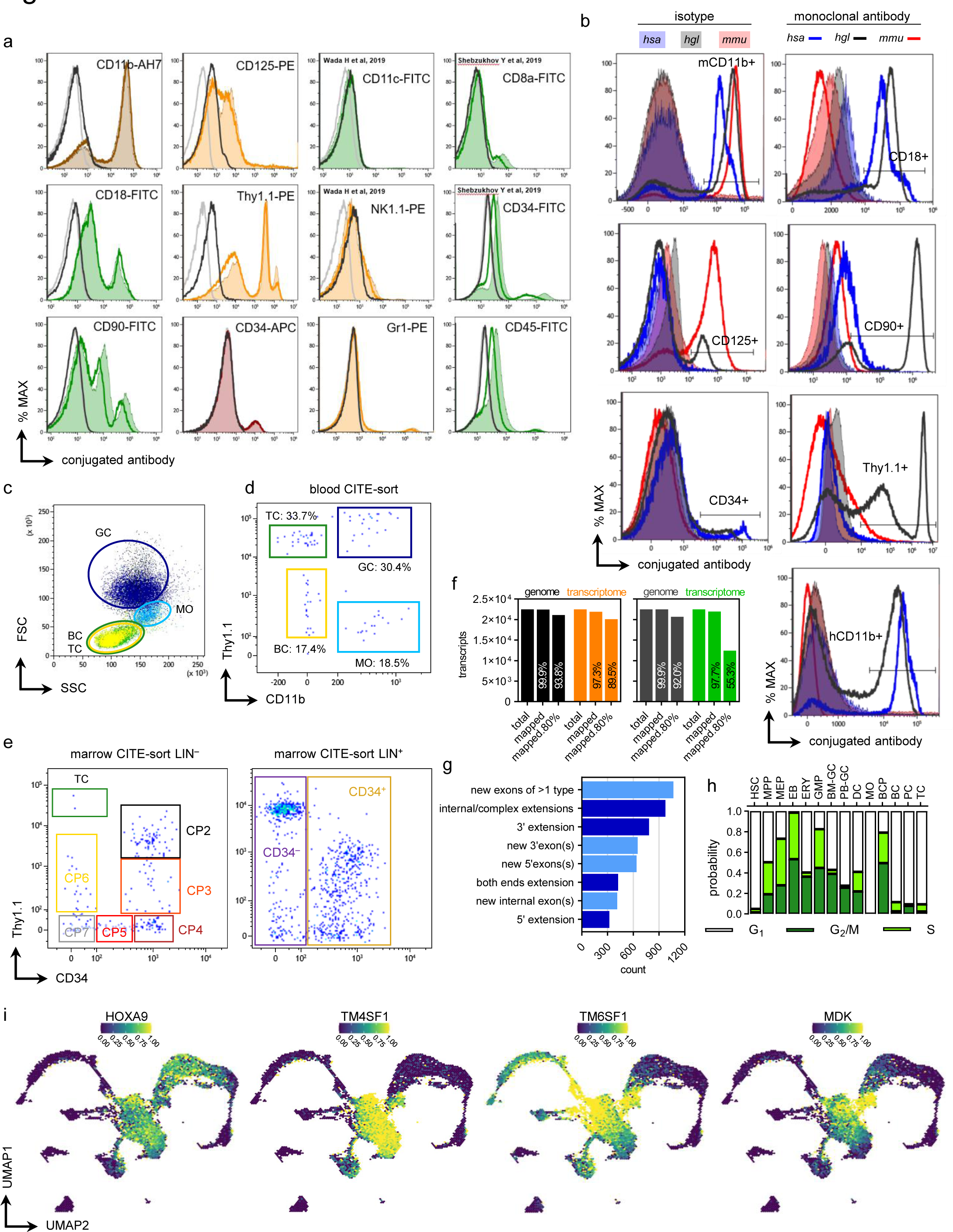
FACS antibody staining pattern validation, transcriptome assembly statistics, sorting input for naked mole-rat scRNA-Seq, literature scRNA-Seq datasets. **a**, Single color flow cytometry histograms of naked mole-rat BM stained for indicated surface markers with off-the-shelf monoclonal FACS antibodies (moAb), conjugates are suffixed as: FITC, fluorescein isothiocyanate; PE, phycoerythrin; APC, allophycocyanin; AH7, APC-Cy7. Grey line, unstained BM; black line, isotype; colored line, Fc Blocker cocktail + moAb; colored Area-Under-Curve (AUC), moAb. Antibodies cross-reactive with naked mole-rat cells described previously are indicated by the respective reference in the top left histogram corner. **b**, Single color flow cytometry histograms of human (*hsa*), mouse (*mmu*) and naked mole-rat (*hgl*) BM stained for indicated surface markers; solid line, Fc Blocker cocktail + isotype; shaded AUC, Fc Blocker cocktail + moAb. **c**, FACS backgating of indicated populations from Figure 1B into side (SSC) and forward (FSC) scatters. Major blood cell type gates for lymphocytes, monocytes and granulocytes were gated and colored as shown in Fig. 1b. **d**, Postsort with 92 viable events of the sorting strategy for PB CITE-Seq, sorting gates referring to Fig. 1b. **e**, Postsort of LIN^−^ BM (left panel) with 242 viable events, sorting gates referring to Figure 2C; LIN^+^ BM (right panel) with 1027 viable events; sorting gates indicated. **f**, FRAMA transcriptome mapped to ENSEMBL genome and transcriptome using cds (left) or full transcript sequence (right). Total, total annotated transcripts; mapped, blastn FRAMA exon contig alignment mapped to genomic coordinates (>95% identity, e-value<1e^-5^); mapped 80%, mapping FRAMA exon contig with >80% mapping coverage (percent of contig length covered by blastn alignments). **g**, Structural differences in FRAMA transcripts with a difference in mapping coverage between genome and ENSEMBL transcriptome of >20%; lightblue, transcripts with new exons; darkblue, transcripts with exon extensions. **h**, *Seurat* cell cycle scoring of sorted naked mole-rat PB and BM (NMR dataset) with clustering from Fig. 1g. **i**, UMAP-based hexbin projection of NMR dataset with clustering from Fig. 1g; scaled expression as probability for each conserved gene. HOXA9 3.6-fold up in HSC, 2.1-fold up in MPP; TM4SF1 6.7-fold up in HSC, 3.9-fold up in MPP; TM6SF1 3.5-fold up in BM-GC, 2.7-fold up in PB-GC, 2.5-fold up in GMP; MDK 4-fold up in MEP, 3.1-fold up in MPP, 2.1-fold up in HSC.

**Fig. S2.**
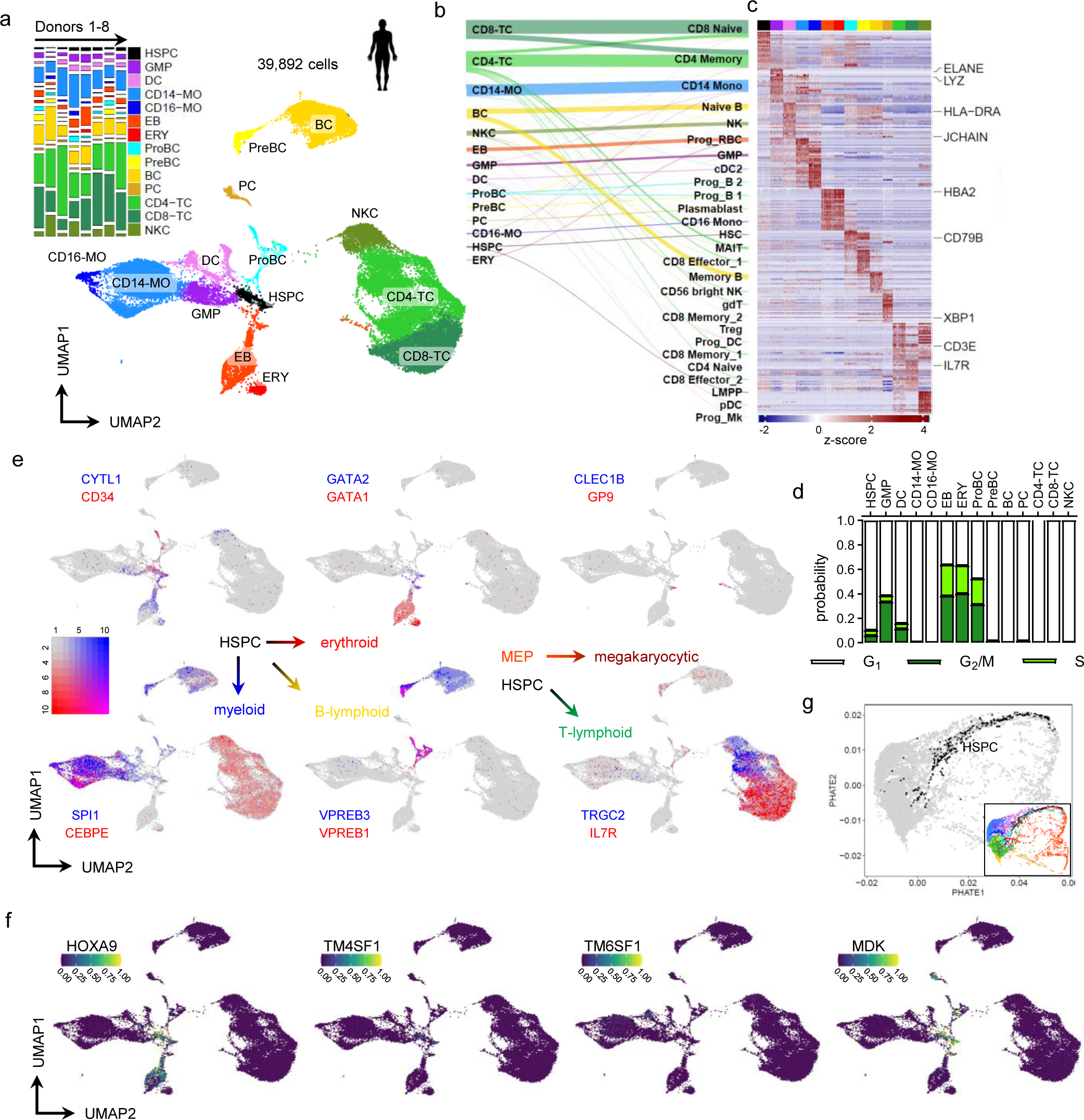
The Human Cell Atlas (**HCA**) dataset as random down-sampled per-donor version distributed via the SeuratData package. **a**, UMAP of Louvain-clustered single-cell transcriptomes, color legend is used throughout this dataset; 4603 differentially expressed genes were used for fGSEA-based cell type annotation. Tile-stack inset reflects relative cluster frequencies [y] and donor library fractions of the dataset [x] as probability. HSPC, hematopoietic stem progenitor cells; CD14-MO, CD14^+^/CD16^lo^ monocytes; CD16-MO, CD14^+^/CD16^hi^ monocytes; ProBC, B progenitor; PreBC, B precursor; CD4-TC, CD4^+^/CD8^−^T cell; CD8-TC, CD4^−^/CD8^+^ T cell; NKC, natural killer cell. **b**, Sankey diagram connecting annotated Louvain clusters [left] with MNN-projected cell identities from a separate human BM CITE-Seq dataset [right] (*87*). **c**, Heatmap showing top 25 overexpressed genes by fold-change of 14 single cell clusters from BM mononuclear cells randomly downsampled to ≤ 500 cells, NMR dataset heatmap marker orthologs are labelled. **d**, *Seurat* cell cycle scoring of HCA dataset. **e**, UMAP-based Blendplots showing pairs conserved lineage markers; gene1 (red, high expression), gene2 (blue, high expression) and co-expressing cells (purple). See scale on the right; expression, scaled UMI counts. **f**,(O) UMAP-based hexbin projection of HCA dataset; scaled expression as probability for each conserved gene. HOXA9 low in HSPC, EB; TM4SF1 very low in HSPC; TM6SF1 not detected; MDK 3.2-fold up in HSPC. NMR dataset: HOXA9 3.6-fold up in HSC, 2.1-fold up in MPP; TM4SF1 6.7-fold up in HSC, 3.9-fold up in MPP; TM6SF1 3.5-fold up in BM-GC, 2.7-fold up in PB-GC, 2.5-fold up in GMP; MDK 4-fold up in MEP, 3.1-fold up in MPP, 2.1-fold up in HSC. **g**, PHATE dimensionality reduction of single-cell transcriptomes, HSPC cluster is highlighted in black; inset depicts model colored by annotation from (J), showing the HSPC cluster linked to erythroid, lymphoid and myeloid branch clusters.

**Fig. S3.**
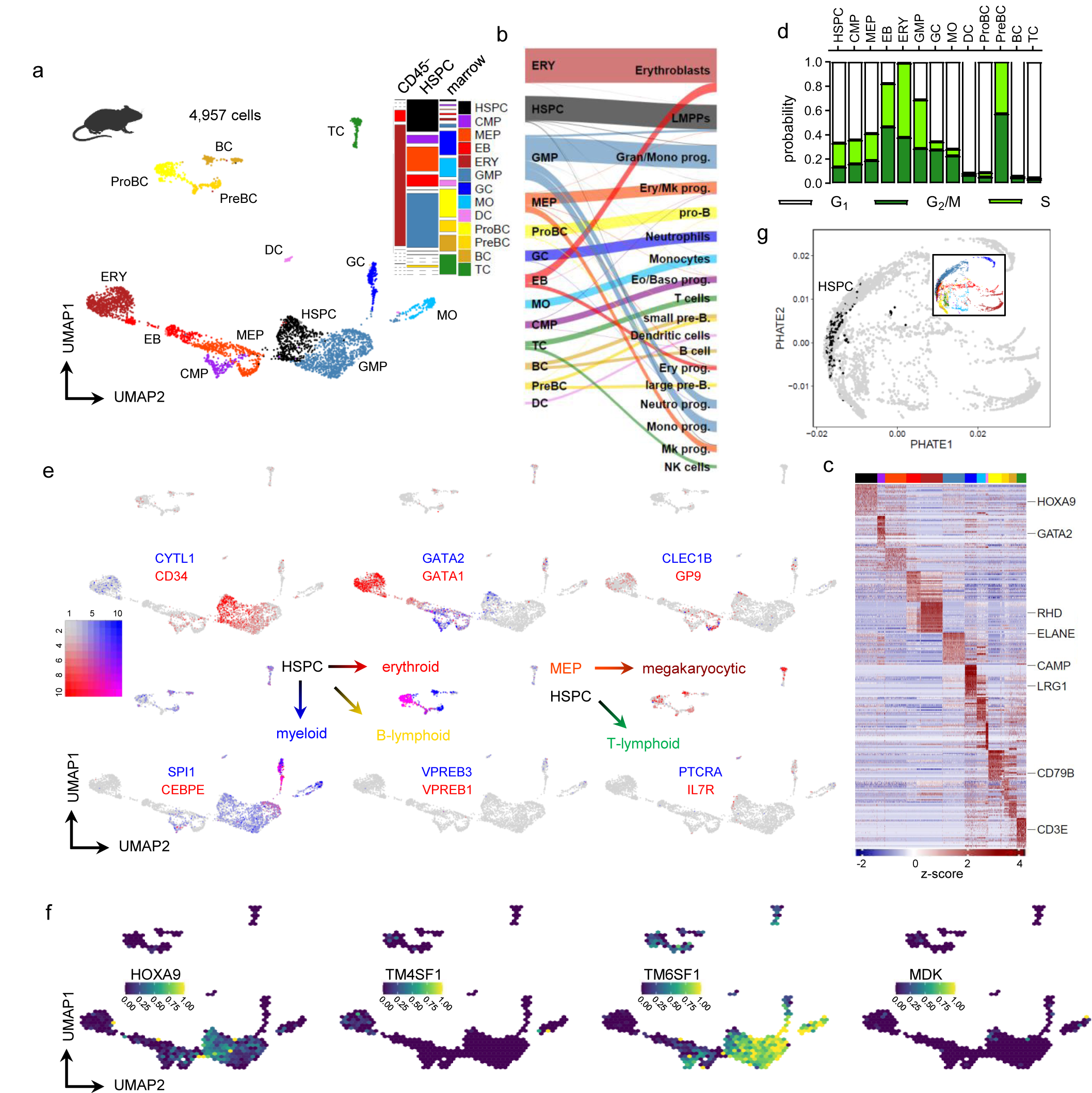
The RNA Magnet (RM) droplet dataset (***85***). **a**, UMAP of Louvain-clustered single-cell transcriptomes, color legend is used throughout this dataset; 1906 differentially expressed genes were used for fGSEA-based cell type annotation. Tile-stack inset reflects relative cluster frequencies [y] and sorted fractions of the dataset [x] as probability. CMP, common myeloid progenitor. **b**, Sankey diagram connecting annotated Louvain clusters [left] with original RNA Magnet annotation [right]. **c**, Heatmap showing top 25 overexpressed genes by fold-change of 13 single cell clusters from sorted marrow cells randomly downsampled to ≤ 500 cells, NMR dataset heatmap marker orthologs are labelled. **d**, *Seurat* cell cycle scoring of TM dataset. **e**, UMAP-based Blendplots showing pairs conserved lineage markers; gene1 (red, high expression), gene2 (blue, high expression) and co-expressing cells (purple). See scale on the right; expression, scaled UMI counts. **f**, UMAP-based hexbin projection of RM dataset; scaled expression as probability for each conserved gene. HOXA9 2.3-fold up in HSPC; TM4SF1, MDK not detected; TM6SF1 high in GMP, GC, MO. **g**, PHATE model of RM dataset, HSPC cluster is highlighted in black; inset depicts color annotation from a, showing the HSPC cluster connected to erythroid, lymphoid and myeloid clusters.

**Fig. S4.**
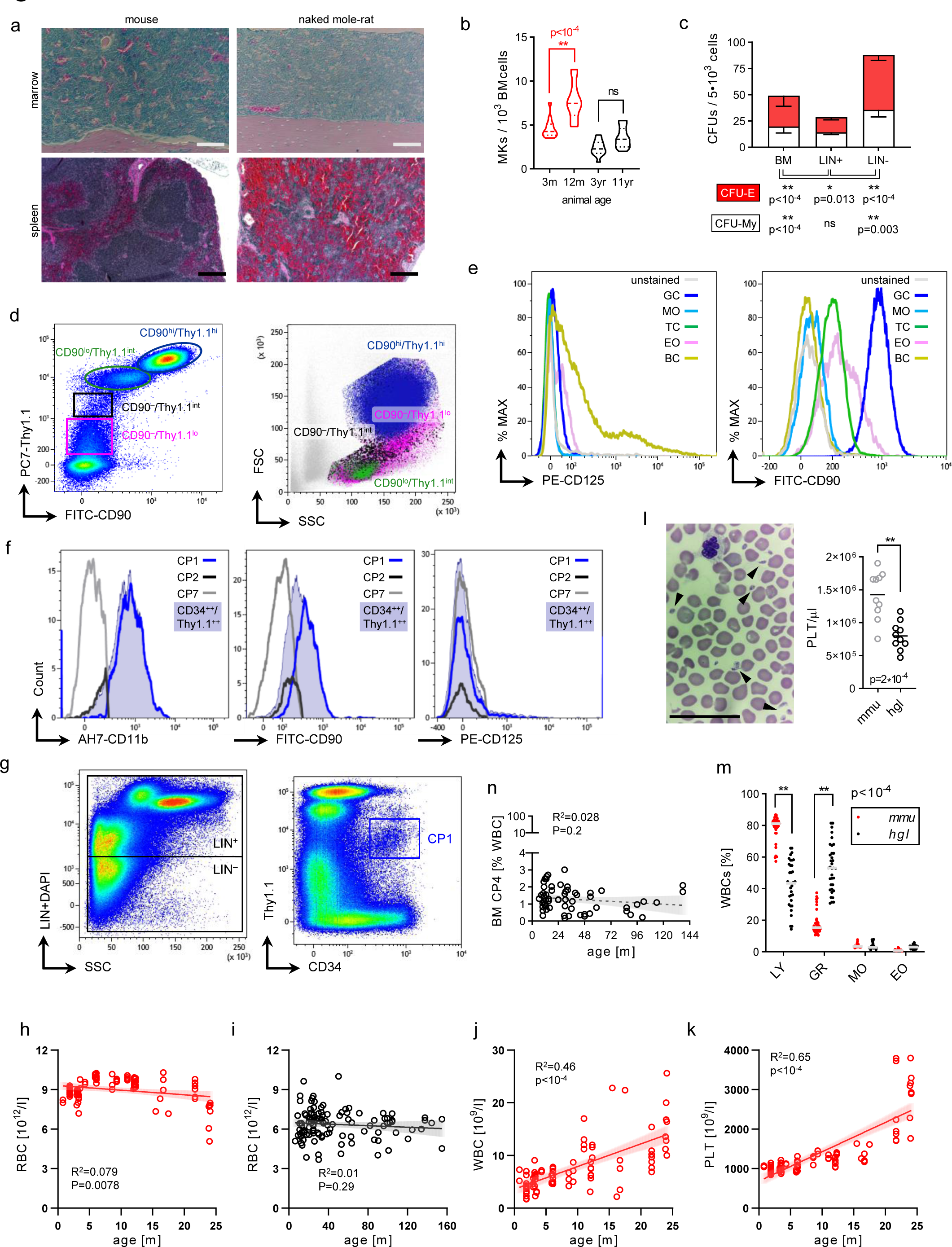
Lineage cocktail (LIN) validation, Hemanalyzer data. **a**, Giemsa staining of femur sections, medullary canal of diaphysis; Scale bar 100µm [top]. Hematoxylin & Eosin staining of spleen sections; Scale bar 200µm [bottom]. **b**, Megakaryocyte (MK) Scoring of BM Giemsa sections; red, mouse; black, naked mole-rat. P-values were determined by One-way ANOVA; n=2. **c**, Colony assay grown at 37°C; CFU-E, colony forming unit erythroid; CFU-My, colony forming unit myeloid. Error bars denote s.d., p-value determined by Sidak’s Two-way ANOVA; n=4. **d**, Viable BM cells stained with CD90 and Thy1.1 [top] give rise to two double positive populations. Backgating into scatter channels [bottom] revealed CD90^hi^/Thy1.1^hi^ cells are restricted to granulocyte scatter properties (darkblue) as were Thy1.1^hi^/CD11b^+^ PB-GCs, while CD90^lo^/Thy1.1^int^ cells appear to be lymphocytes (green), as seen for Thy1.1^int^/CD11b^−^ TCs in blood. CD90^−^/Thy1.1^int^ and CD90^−^/Thy1.1^lo^ cells contain naked mole-rat HSPCs and have heterogeneous size and low granularity, compared to human HSPCs predominantly sized between lymphocytic and monocytic leukocyte types (*88*). FITC, fluorescein isothiocyanate; PC7, PE-Cy7; SSC, side scatter; FSC, forward scatter. **e**, Histograms of CD125 [left] and CD90 [right] of PB leukocytes with cell type gates from Fig. 1b; % MAX, scales the maximum of all datasets at the same level. CD125 is exclusively found a BC subset, GCs are CD90^hi^ and Eos, TCs are CD90^lo^, BCs, MOs are CD90^−^. **f**, Signal intensities of CD11b [left], CD90 [middle] and CD125 [right] of indicated BM cell fractions; Grey line, CP7; black line, CP2/LTC; blue line, CP1; viable CD34^hi^/Thy1.1^int^ not LIN gated, steelblue AUC. FITC, fluorescein isothiocyanate; PE, phycoerythrin; AH7, APC-Cy7. **g**, Sorting strategy for spleen cells with lineage (LIN) depletion [left] and gating of LIN^+^ CP1 [right]; Note the prominent LIN^dim^ population as a spleen-specific staining pattern defining the upper limit for the LIN^−^ boundary. We set the LIN^−^ gate at the transition towards LIN^+^/SSC^low^ cells similar to BM (Fig. 2a). In mice the frequency of repopulating, self-renewing stem cells is ∼10-fold lower in spleen compared to BM (*89*). Median BM LTC/CP2 are 0.151%, median spleen LTC 0.038%, ∼4-fold lower in spleen (Fig. 2c-f). Hemanalyzer volumetric RBC numbers across animal age for **h**, C57BL/6 mice (n=88) or **i**, naked mole-rats (n=115). R^2^, Pearson correlation coefficient; p-values were determined by conventional linear regression fitting both slope and intercept. Volumetric **j**, white blood cell (WBC) and **k**, platelet (PLT) numbers for C57BL/6 mice. R^2^, coefficient of determination; p-values were determined by conventional linear regression fitting both slope and intercept; n=88. **l**, Wright staining of naked mole-rat PB film [left], arrows indicate scored platelets; Scale bar 50µm. Scoring of PB films [right], p-values were determined by unpaired Student’s t-test; n=10. **m**, Hemanalyzer WBC subset frequencies (% of total leukocytes); p-values were obtained from Sidak’s two-way ANOVA. Mouse, n=45; naked mole-rat, n=36. **n**, BM CP4 frequencies across age; n=60, linear regression with 95% CI as trend line; p<0.05, significance.

**Fig. S5.**
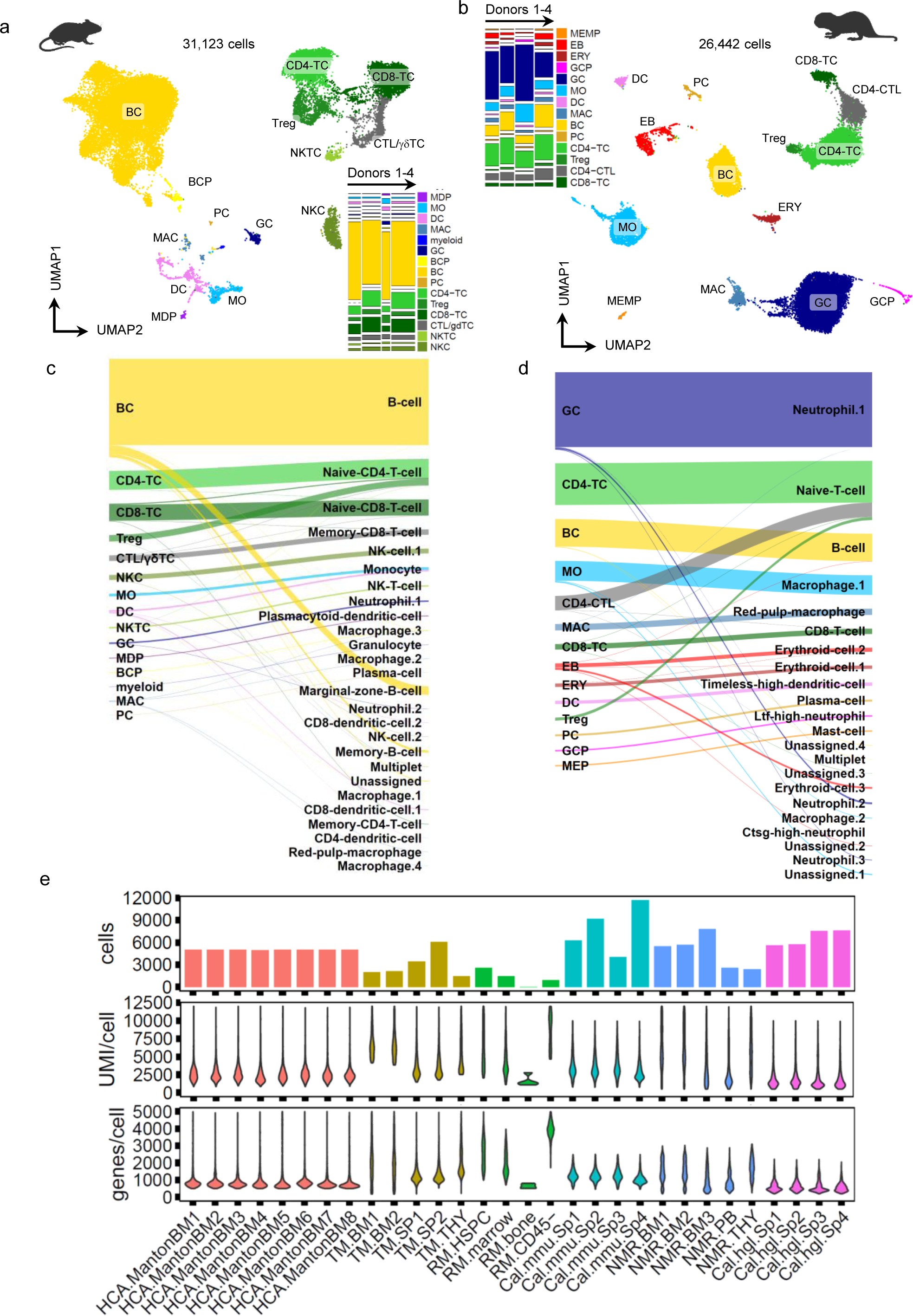
The Calico droplet dataset (***26***). UMAP of Louvain-clustered **a**, mouse or **b**, naked mole-rat whole spleen single-cell transcriptomes, color legend is used throughout this dataset; 1906 differentially expressed genes were used for fGSEA-based cell type annotation. Tile-stack inset reflects relative cluster frequencies [y] and donor fractions of the dataset [x] as probability. MDP, monocytic dendritic progenitor. Sankey diagram connecting annotated Louvain clusters [left] with original Calico annotation [right] for **c**, mouse or **d**, naked mole-rat. **e**, Quality metrics for all scRNA-Seq datasets from Fig. 1, Fig. S2, 3, 5. Top, number of single cells per demultiplexed and filtered library; Middle, UMI counts per cell for each library; Bottom, detected genes per cell for each library. BM, bone marrow; SP, spleen; THY, thymus; *mmu*, mouse; *hgl*, naked mole-rat.

**Fig. S6.**
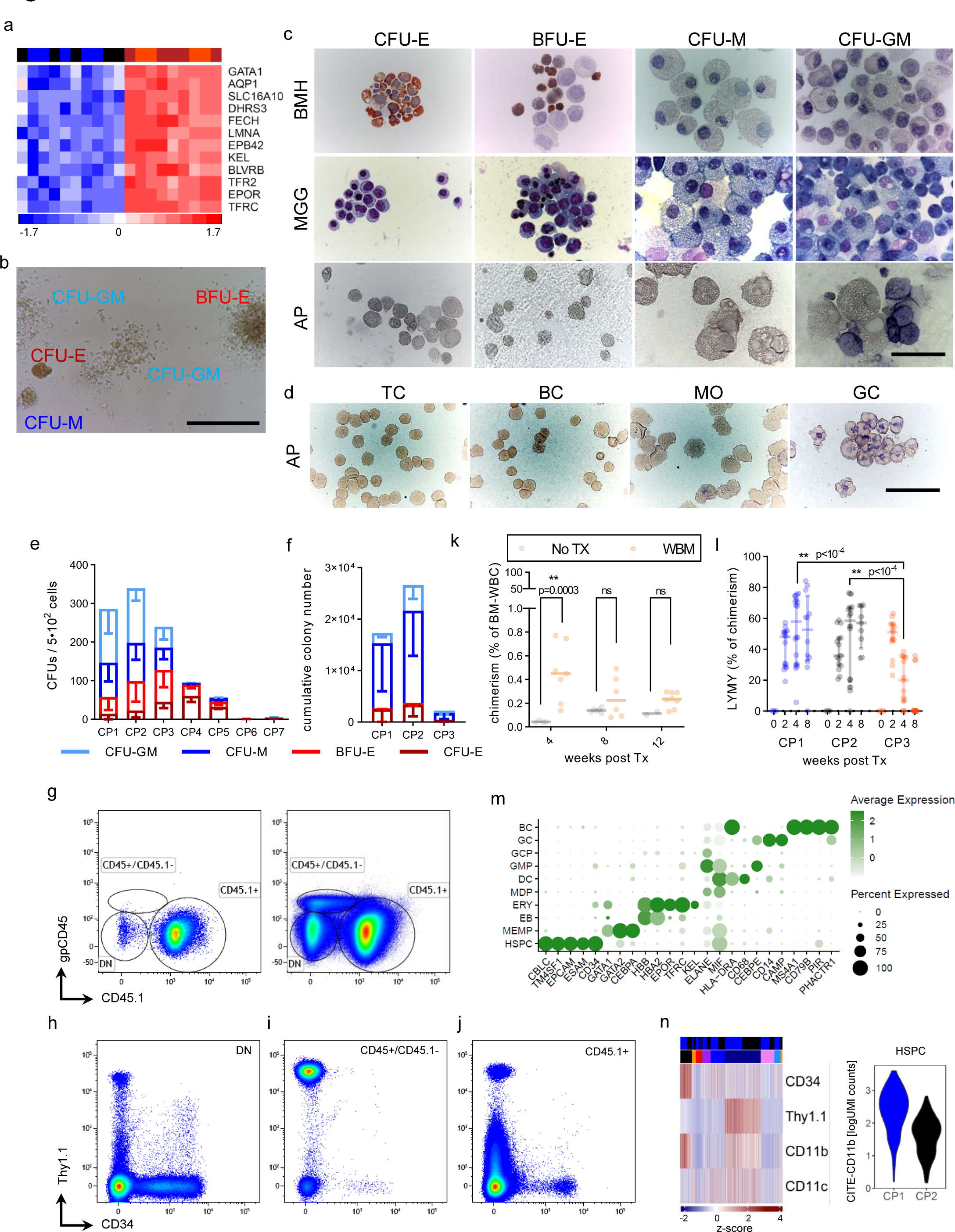
BM CP1-4 bulk RNA-Seq, Colony assays, Xenografts. **a**, Top color bar reflects population clustering by Euclidean distance using top 12 MEP/erythroid leading edge genes; color key as in Fig. 3a. Genes were obtained by ssGSEA of combining CP1/2 (n=10) versus CP3/4 (n=10) populations and performing differential gene expression (DGE) testing, see also Table S4. **b**, Representative image of colony morphologies grown at 32°C. BFU-E, burst forming unit erythroid; CFU-E, colony-forming unit erythroid; CFU-M, colony forming unit macrophage; CFU-GM, colony-forming unit granulocyte-monocyte; Scale bar 250µm. **c**, Benzidine (top row, BMH), May-Grünwald-Giemsa (2^nd^ from top, MGG) and Alkaline Phosphatase (3^rd^ from top, AP) stainings of picked colonies; scale bar 50µm. Erythroid CFU and BFU cells contained hemoglobin, the latter at lower frequency than the more mature CFU-E. CFU-M exclusively consisted of large cells with colorless vesicles and a high cytoplasmic/nuclear ratio, while CFU-GM (granulocyte/monocyte) also comprised AP^+^ cells. **d**, Alkaline Phosphatase (AP) staining of sorted PB cells (bottom row); scale bar 50µm. Naked mole-rat PB-GCs are AP^+^, other WBC subsets AP^−^. **e**, Colony assay (n=5) or **f**, Replated assays (n=3) of sorted naked mole-rat BM CPs, grown at 32°C. Error bars denote s.d., p-value determined by Sidak’s Two-way ANOVA. **g**, Untransplanted [right] and naked mole-rat CP2-transplanted [left] NSGS host BM stained with guinea-pig CD45 and CD45.1; CP2 xenograft at week 4; DN, double negative. **h**, DN fraction contains most CD34^+^ and Thy1.1^+^/CD34^+^ naked mole-rat cells and a Thy1.1^−^/CD34^−^ population potentially containing xenogenic CP7; **i**, CD45^+^/CD45.1^−^ fraction contains most naked mole-rat Thy1.1^hi^/CD34^−^ BM-GCs; **j**, CD45.1^+^ fraction features an abundant Thy1.1^lo^/CD34^−^ population resembling CP6 in naked mole-rat BM and spleen. **k**, Recipient chimerism in BM; WBM, naked mole-rat whole bone marrow. **l**, Kinetics of engraftment proportions for LYMY gate from Fig. 4d; p-value determined by Tukey’s Two-way ANOVA. **m**, Top differentially expressed cell type markers of integrated CP1/CP2 xenograft scRNA-Seq. **n**, CITE-signals for the complete integrated xenograft scRNA-Seq dataset [right]. Graft color bar: CP1, blue; CP2, black. Cell type annotation bar colors refer to Fig. 4l. CITE-CD11b levels per cell [right] for the HSPC cluster from the integrated dataset.

**Fig. S7.**
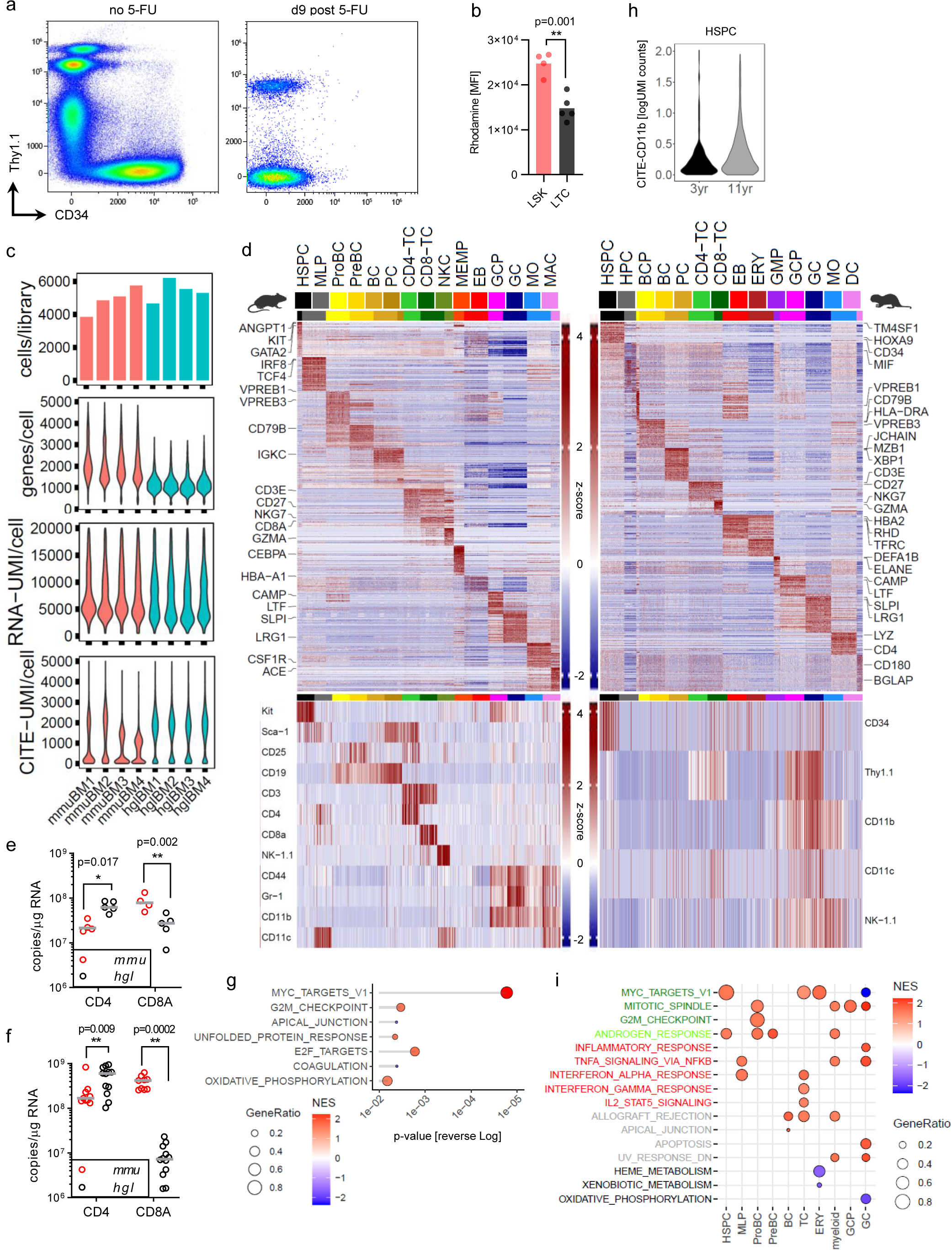
Unfractionated BM scRNA-Seq QA, hierachical clustering, CD4:CD8 qPCR, age groups, GSEA. **a**, Naked mole-rat spleen cells untreated or treated with 5-FU. A marrow Thy1.1^int^/CD34^hi^ HSPC population (Figure 3M) and an expanded splenic Thy1.1^−^/CD34^lo/hi^ erythroid compartment are clearly visible in untreated animals (1-3 year old). By contrast, 5-FU eliminates the entire erythroid lineage in both organs and reduces marrow HSPCs to Thy1.1^hi^/CD34^hi^ myeloid progenitors. **b**, Mean Fluorescence Intensity (MFI) of Rhodamine efflux measurements in mouse (n=4) and naked mole-rat (n=5) BM. p-value determined by unpaired Student’s t-test. **c**, Quality metrics for BM scRNA-Seq datasets from Fig. 5. Top, number of single cells per demultiplexed and filtered library; 2^nd^ from top, detected genes per cell; 3^rd^ from top, mRNA UMI counts per cell; Bottom, CITE UMI counts per cell for each library. BM, bone marrow; *mmu*, mouse; *hgl*, naked mole-rat. No cell enrichment procedure was applied prior to sequencing, and RBC removal through osmolysis rather than Percoll-based methods was used to capture the native marrow WBC content from both species. **d**, Top heatmaps show the top 25 cell type specific markers for mouse [left] and naked mole-rat [right] BM randomly downsampled to ≤ 500 cells/cluster, canonical cell type markers from the literature are labelled. Bottom heatmaps show the cell type specific CITE features for mouse [left] and naked mole-rat [right] BM randomly downsampled to ≤ 100 cells/cluster. Fold-change cut-off 2, p-value threshold 0.05. **e**, Absolute copy number determination by qPCR for CD4 and CD8A transcripts in sorted PB-TCs from mouse (Cd11b^−^ /Gr-1^−^/Cd19^−^/Cd3e^+^; n=4) and naked mole-rat (CD11b^−^/Thy1.1^int^; n=5). P-values derived from Sidak’s two-way ANOVA. **f**, Absolute qPCR in whole cervical lymph nodes from mouse (n=9) or naked mole-rat (n=12). P-values derived from Sidak’s two-way ANOVA. **g**, GSEA with the MSigDB hallmark geneset collection of differentially expressed genes across each individual cluster (n=760) between 12 month vs 3 month old mice; p-value threshold 0.05. NES, normalized enrichment score; GeneRatio, (signature ∩ term) / (signature ∩ all terms). **h**, CITE-CD11b levels between 3 year vs 11 year HSPCs within naked mole-rat BM. **i**, GSEA with the MSigDB hallmark geneset collection of differentially expressed genes across each individual cluster (n=7206) between SCTransform-integrated mouse vs naked mole-rat BM; FDR q-value threshold 0.05. NES, normalized enrichment score; GeneRatio, (signature ∩ term) / (signature ∩ all terms).

**Fig. S8.**
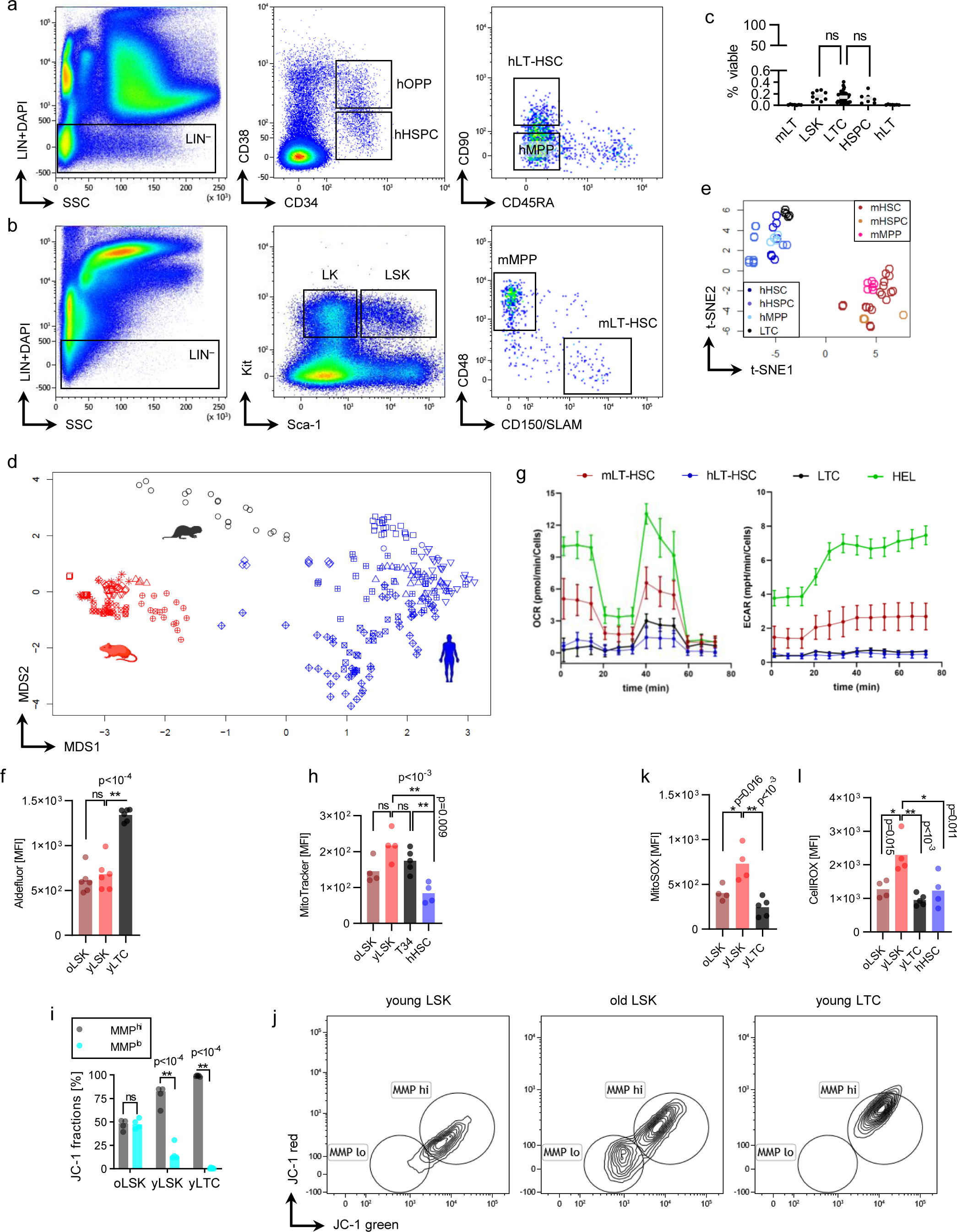
Cross-species bulk RNA-Seq, Seahorse, MMP. **a**, Representative gating strategy for human BM with lineage cocktail (LIN; CD3/CD14/CD16/CD19/CD20/CD56) to select LIN^−^ cells [left]; gating LIN^−^/CD38^lo^/CD34^+^ human HSPCs or LIN^−^/CD38^+^/CD34^+^ human oligopotent progenitors (hOPP) [middle]; gating LIN^−^/CD38^lo^/CD34^+^/CD90^+^/CD45RA^−^ long term LT-HSCs and LIN^−^/CD38^lo^/CD34^+^/CD90^−^ /CD45RA^−^ multipotent progenitors (hMPP) [right]. **b**, Representative gating strategy for mouse BM with lineage cocktail (LIN; Cd3e/Gr-1/Cd11b/B220/Ter-119) to select LIN^−^ cells [left]; gating LIN^−^/Sca-1^+^/Kit^+^ (LSK) and LIN^−^/Sca-1^−^/Kit^+^ (LK) [middle]; LIN gate set to 10% viable cells in a biplot analog to Figure 2A. **c**, Frequencies of BM HSC compartments between species. Mouse mLT-HSC (n=7); mouse LSK (n=10); naked mole-rat LIN^−^/Thy1.1^int^/CD34^+^, LTC (n=47); human HSPC (n=7); hLT-HSC (n=7); p-value determined by Dunnett’s One-way ANOVA. **d**, Unsupervised clustering by multi-dimensional scaling (MDS) of 299 bulk RNA-Seq samples. Mouse, red (n=100); Human, blue (n=179); Naked mole-rat, black (n=20). See Table S6 for sample metadata. **e**, Unsupervised clustering by t-distributed stochastic neighborhood embedding (t-SNE) of Primitive stem and progenitors. hHSC, LIN^−^/CD34^+^/CD38^lo^/CD90^+^/CD45RA^−^ (n=7); hHSPC, LIN^−^/CD34^+^/CD38^lo^ (n=11); hMPP, LIN^−^/CD34^+^/CD38^lo^/CD90^−^/CD45RA^−^ (n=4); mHSC, LIN^−^/Sca-1^+^/Kit^+^/CD150^+^/CD48^−^ (n=25); mMPP, LIN^−^/Sca-1^+^/Kit^+^/CD150^−^/CD48^+^ (n=6); LTC, LIN^−^/Thy1.1^int^/CD34^+^ (n=5). **f**, Mean Fluorescence Intensity (MFI) of Aldefluor measurements (n=6); p-value determined by Tukey’s One-way ANOVA. **g**, Seahorse XF Cell Mito Stress Test profiles for HSPCs. HEL, human erythroleukemia cell line; used as positive control on all assay plate run with different primary cells. **h**, MFI of Mitotracker Red staining in mouse (n=4), human (n=4) and naked mole-rat BM (n=5); p-value determined by Tukey’s One-way ANOVA. **i**, Proportions of JC-1 mitochondrial membrane potential (MMP) populations mouse (n=4; yLSK, 3 month old; oLSK, 24 month) and naked mole-rat (n=5) BM. p-value determined by Sidak’s Two-way ANOVA. j, Merged (composite analysis with all individual samples concatenated into one dataset) JC-1 fluorescence biplots for young LSK (3m old, yLSK) [right], old LSK (24m old, oLSK) [middle], young LTC (2-3yr old). MFI of **k**, MitoSOX or **l**, CellROX from mouse (n=4), human (n=4) or naked mole-rat BM (n=5); p-value determined by Tukey’s One-way ANOVA.

**Fig. S9.**
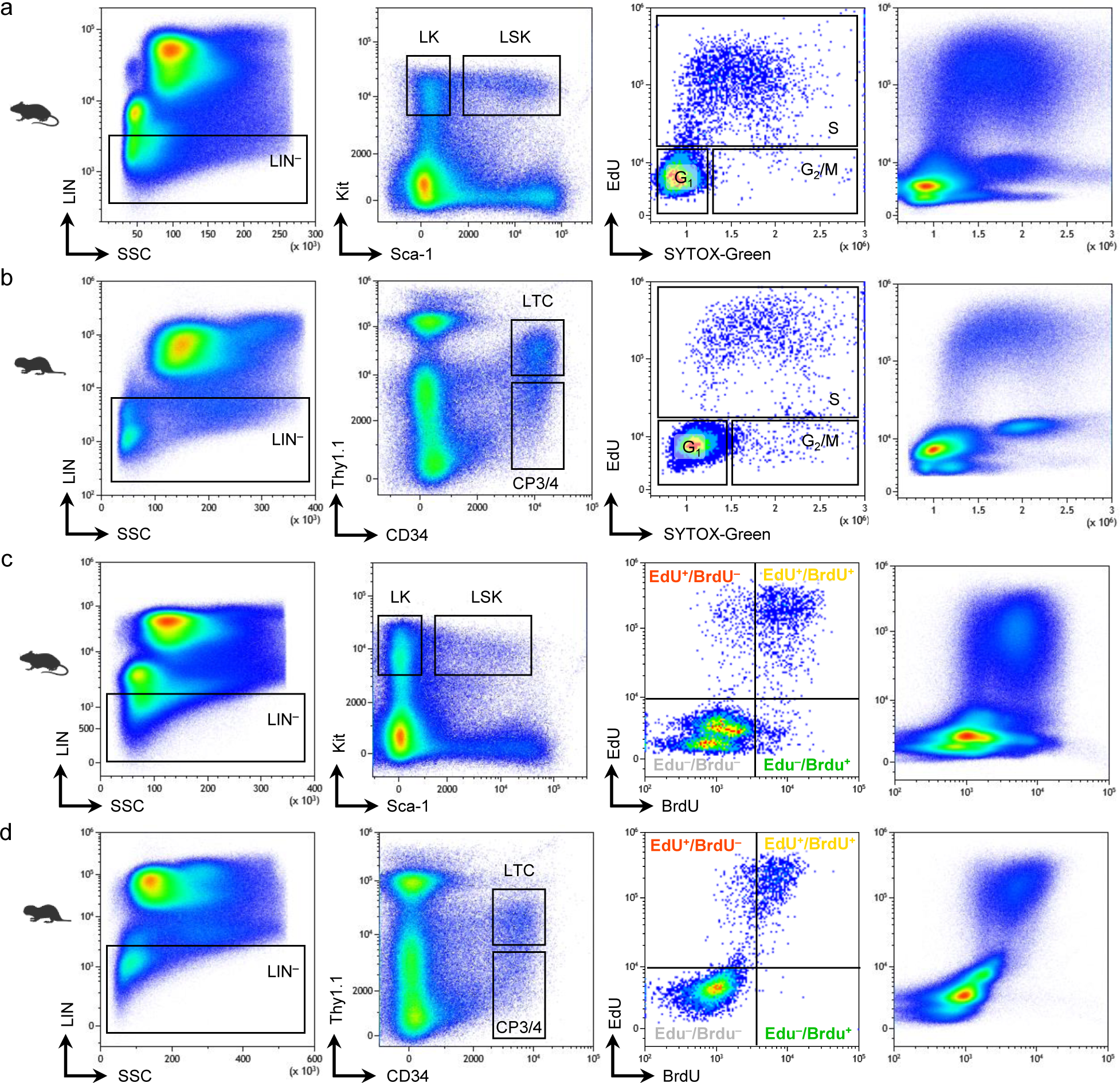
Dual-Pulse FACS gating. For each species individual sample data was combined to one merged dataset to illustrate gating and population patterns; mouse, n=7; naked mole-rat, n=5. Gating path for EdU/DNA-content cell cycle analysis of **a**, mouse or **b**, naked mole-rat BM. Lineage depletion [left] used to enrich for HSPCs and progenitors [2^nd^ from left]; CP3/4, compound gate of CP3 and CP4, see Fig. 3c. Specific gates for each cell cycle according to EdU-label and DNA content on mouse LSK or naked mole-rat LTC [3^rd^ from left]. EdU over DNA-content for all viable cells of the complete mouse or naked mole-rat BM dataset, respectively [left]; Note that naked mole-rats featured less S-phasing cells in whole marrow than mice. Gating path for EdU/BrdU analysis of **c**, mouse or **d**, naked mole-rat BM. Lineage depletion [left] used to enrich for HSPCs and progenitors [2^nd^ from left]. Specific gates for each fluorescent fraction on mouse LSK or naked mole-rat LTC [3^rd^ from left]. EdU over BrdU for all viable cells of the complete mouse or naked mole-rat BM dataset, respectively [left]; Note that naked mole-rat marrow lacks a EdU^−^/BrdU^+^ population.

